# Molecular basis of promiscuous chemokine-engagement by the Duffy antigen receptor

**DOI:** 10.64898/2026.03.09.710718

**Authors:** Manisankar Ganguly, Yuma Matsuzaki, Nabarun Roy, Divyanshu Tiwari, Sameer Kulkarni, Annu Dalal, Manish K. Yadav, Nilanjana Banerjee, Sudha Mishra, Kazuhiro Sawada, Kana M. Hashimoto, Kohei Yamaguchi, Martin J. Stone, Richard J. Payne, Andy Chevigne, Fumiya K. Sano, Ramanuj Banerjee, Osamu Nureki, Arun K. Shukla

**Author notes:** Joint 1st author.

## Abstract

The Duffy blood group antigen, encoded by a seven transmembrane protein known as Duffy Antigen Receptor for Chemokines (DARC), or Atypical Chemokine Receptor 1 (ACKR1), serves as a key receptor for the malarial parasite, *Plasmodium vivax,* on erythrocytes. DARC exhibits remarkable functional divergence compared to prototypical chemokine receptors and other G protein-coupled receptors (GPCRs) as it does not engage canonical signal-transducers such as G-proteins, GPCR regulatory kinases (GRKs), and β-arrestins. DARC is a highly promiscuous receptor interacting with several homeostatic and inflammatory C-C and C-X-C subtype chemokines, and similar to other chemokine receptors, these interactions are modulated by post-translational tyrosine sulfation in the N-terminus. A single nucleotide polymorphism in DARC leading to Gly^42^Asp substitution forms the basis for Fy^a^ vs. Fy^b^ Duffy blood group antigen classification with differential impact on *Plasmodium vivax* infection and cancer progression. However, the molecular basis of promiscuous chemokine-binding by DARC and the modulation by receptor sulfation and Fy^a^/Fy^b^ allelic variation remains unclear. Here, we design a sortase-mediated chemical-ligation strategy to generate purified DARC with naturally-occurring tyrosine sulfation at the N-terminus, and determine high-resolution cryo-EM structures in complex with a C-C type chemokine, CCL7, and a C-X-C type chemokine, CXCL8. We observe that similar to CCL7, CXCL8 engages with DARC primarily through the N-terminus of the receptor, which is in stark contrast with the two-site binding mechanism displayed by other chemokine receptors. Interestingly, the E-L-R motif in CXCL8 engages a pseudo-R-D motif in DARC leading to superficial engagement unlike CXCR2, a prototypical chemokine receptor sharing the same agonist. Moreover, tyrosine sulfation and Fy^b^ allelic variation leads to a repositioning of the receptor N-terminus on the core domain of CXCL8, resulting in distinct interaction network imparting greater binding affinity. Taken together, our study presents novel insights into promiscuous chemokine-binding to DARC, tyrosine sulfation-mediated fine-tuning of chemokine engagement, and a generalizable sortase-mediated chemical-ligation platform applicable to other chemokine receptors.

## Introduction

The Duffy antigen receptor for chemokines (DARC), also referred to as Atypical Chemokine Receptor subtype 1 (ACKR1), is a seven transmembrane receptor (7TMR) expressed primarily on erythrocytes, epithelial and endothelial cells, and airway smooth muscle cells ^1,2^. It binds multiple chemokines across the C-C and C-X-C subtypes, but does not transduce downstream signaling via canonical pathways utilized by prototypical chemokine receptors and other G protein-coupled receptors (GPCRs)^3–8^. Consequently, DARC is classified as a member of the Atypical Chemokine Receptors (ACKRs) subfamily, although unlike other ACKRs, it fails to elicit any measurable β-arrestin (βarr) recruitment in transfected cells^9^. Interestingly, DARC also serves as the key receptor on the erythrocytes for the entry of the malarial parasite, *Plasmodium vivax*, by interacting with the Duffy binding protein expressed at the surface of the parasite^10–14^. Moreover, when expressed at the endothelial cells, it serves as the key attachment point for the binding and oligomerization of the pore forming toxins (PFTs) secreted by pathogenic bacteria such as *Staphylococcus aureus*^15–18^. More broadly, DARC is proposed to sequester chemokines in the physiological context, thereby modulating the chemokine levels and gradients with direct implications for cancer onset and progression^19–21^ as well as immune cell trafficking in airway inflammation and hyperresponsiveness^22,23^. Therefore, a more detailed understanding of the structure, function, and ligand-recognition by DARC is essential to elucidate a deeper mechanistic framework of host-pathogen interaction with a significant potential to inform novel drug discovery.

We have recently presented a comprehensive characterization of DARC using cellular assays to demonstrate a complete lack of canonical transducer-coupling such as G-proteins, GRKs, and βarrs^9,24–26^. In addition, we have also determined a cryo-EM structure of DARC in complex with CCL7, a C-C subtype chemokine, elucidating the overall binding modality and receptor architecture^9^. Still however, several fundamental questions remain unanswered related to promiscuous chemokine-engagement by DARC, and their fine-tuning by post-translational modifications such as N-terminal tyrosine sulfation. For example, how does DARC interact with both C-C and C-X-C type chemokines, unlike the strong selectivity for either C-C or C-X-C chemokines encoded by other chemokine receptors? We have previously demonstrated that a broadly conserved sequence motif in the C-X-C type chemokines (i.e. E-L-R) engages a structural motif in CXCR2 (i.e. R-D) to orchestrate promiscuous binding of C-X-C chemokines at this receptor^27^. At the same time, this interaction of the sequence motif in the chemokines with the structural motif in the receptor also imparts selectivity against C-C chemokines and other non-CXCR2-binding C-X-C chemokines, which lack the E-L-R motif^27,28^. Similar to other chemokine receptors, DARC also harbors two distinct tyrosine sulfation sites i.e. Tyr^30^ and Tyr^41^, and this post-translation modifications appear to modulate chemokine binding and affinity^29,30^. However, it is not clear how exactly the presence of tyrosine sulfation modulates the interaction of chemokines with DARC at the molecular level. Considering that nearly every chemokine receptor harbors putative tyrosine sulfation sites in their N-terminus (**Figure 1A**), and that previous studies have implicated this modification in chemokine binding^31–37^, deciphering the molecular mechanism is likely to have generalizable implications for the chemokine-chemokine receptor system. Finally, the structural mechanism imparting functional divergence on DARC compared to other chemokine receptors and GPCRs, especially in terms of the overall 7TM fold and cytoplasmic surface that precludes the engagement of canonical transducers also remains to be fully understood.

**Figure 1.**
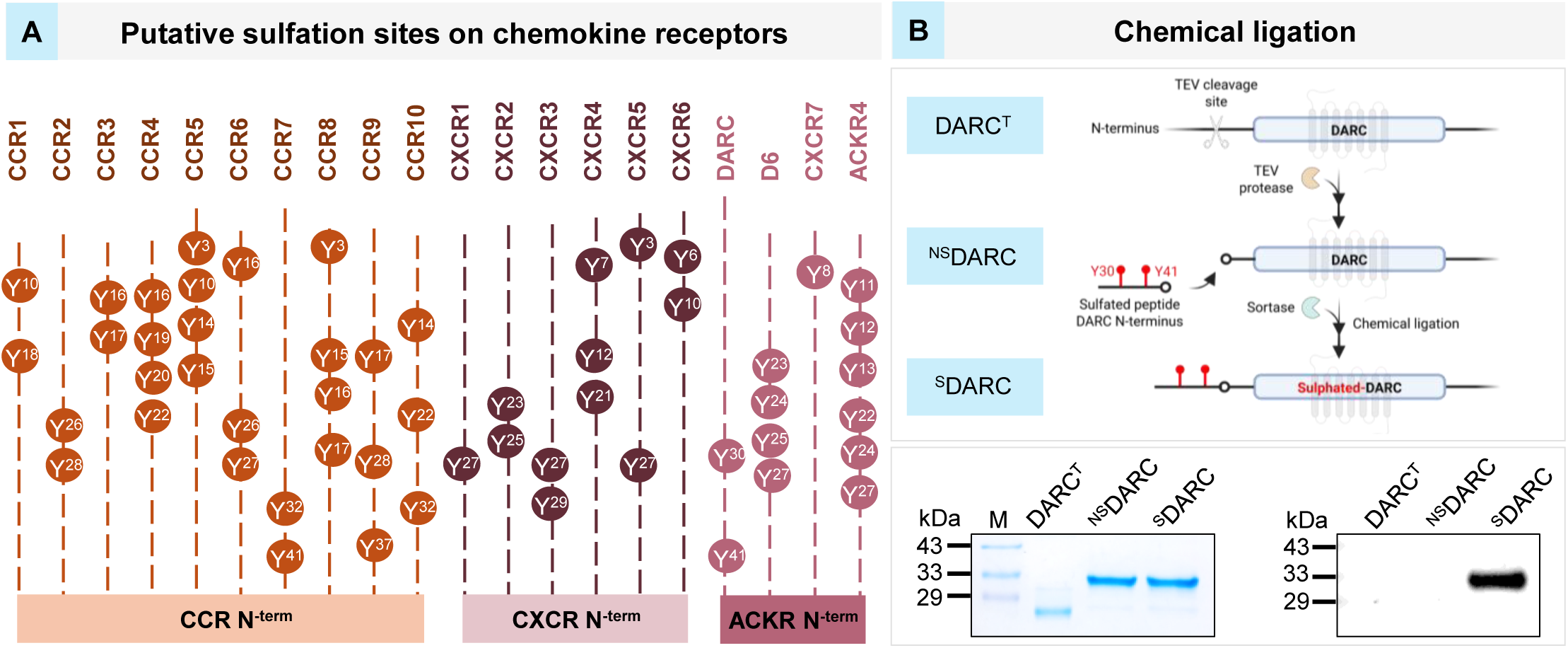
A chemical-ligation strategy to generate ^Sulfo^DARC. **(A)** A schematic representation of putative tyrosine sulfation sites in the N-terminus of chemokine receptors. **(B)** A schematic depicting the experimental strategy to generate uniformly sulfated DARC (i.e. ^Sulfo^DARC) using sortase-mediated chemical-ligation approach, followed by, visualization of ^Sulfo^DARC on Western blot using anti-sulfo-Tyrosine antibody (DARC^T^ = N^term^ cleaved DARC; ^NS^DARC = non-sulfated N^term^ peptide ligated to DARC^T; S^DARC = sulfated N^term^ peptide ligated to DARC). The N^term^ peptide used here (i.e. Gln^19^-Leu^45^) corresponds to the Fy^b^ variant of DARC (i.e. D^42^).

Here, we present a generalizable strategy based on the use of sortase-mediated chemical-ligation to obtain N-terminally-sulfated DARC, and a series of cryo-EM structures to elucidate the molecular details of promiscuous chemokine-binding and fine-tuning by tyrosine sulfation. We also elucidate the impact of Gly^42^Asp allelic variation in DARC, which forms the basis for Fy^a^ vs. Fy^b^ classification of Duffy blood group, on chemokine engagement. These findings provide fundamental and broadly applicable insights into chemokine-chemokine receptor interactions that could be used to inform novel therapeutic approaches. The sortase-mediated chemical-ligation strategy developed for installation of tyrosine sulfation as part of this work serves as a general method for the generation of N-terminally modified receptors and raises up the possibility of precisely assessing the effect of tyrosine sulfation on binding of an array of chemokines to their cognate receptors.

## Results

### A chemical-ligation strategy to generate ^Sulfo^DARC

In this study, we set out to address three key questions pertaining to chemokine engagement with DARC. First, how does DARC recognize both C-C and C-X-C type chemokines? Second, how does tyrosine sulfation in the N-terminus of DARC impact chemokine binding? Third, how does Fy^a^/Fy^b^ allelic variation in DARC fine-tune chemokine recognition? In the previously determined cryo-EM structures of the chemokine-chemokine receptor complexes including CCL7-bound DARC, a discernible density corresponding to tyrosine sulfation is not resolved, presumably due to inefficient tyrosine sulfation in *Sf*9 cells^38–40^. Therefore, we first designed a chemical-ligation strategy to obtain uniform sulfation at Tyr^30^ and Tyr^41^ in the N-terminus of DARC (**Figure 1B**). We engineered a construct of DARC harboring a TEV protease cleavage site in the N-terminus at position Leu^45^, and purified the receptor using baculovirus-mediated expression in *Sf*9 cells followed by anti-FLAG antibody affinity chromatography. Upon TEV cleavage of purified DARC, we obtained N-terminally truncated receptor with a tri-Glycine acceptor sequence (i.e. Gly-Gly-Gly-Glu^46^-Ser^336^) for sortase-based chemical-ligation. The truncated receptor was subsequently incubated with a synthetic peptide corresponding to the N-terminus of DARC (Gln^19^-Leu^45^) harboring sulfated Tyr^30^ and Tyr^41^, and a sortase ligation sequence (LPETGAA) at the carboxyl-terminus, in presence of the sortase enzyme. The ligated receptor bearing pre-defined and uniform tyrosine sulfation at Tyr^30^ and Tyr^41^ (we refer to this as ^Sulfo^DARC herein) was subsequently purified using size-exclusion chromatography and validated using Western blot with anti-Sulfo-Tyr antibody. ^Sulfo^DARC exhibits robust binding to CCL7 and CXCL8, and was obtained in suitable purity for structural analysis.

### Structure determination of chemokine-bound ^Sulfo^DARC

Next, we determined the cryo-EM structures of CCL7-^Sulfo^DARC^G42^ and CXCL8-^Sulfo^DARC^G42^ complexes at global resolutions of 3.3Å and 3.7Å, respectively (**Figure 2A-B and Supplementary Figure S1-2**). In addition, we also generated an Fy^b^ version of ^Non-sulfo^DARC and ^Sulfo^DARC by ligating N-terminal peptides harboring Gly^42^Asp mutation without and with sulfation, respectively, and determined their structures in complex with CXCL8 at overall resolutions of 3.9Å and 3.5Å, respectively (**Figure 2A-B and Supplementary Figure S1-2**). As sortase ligation results in the insertion of a non-natural seven amino acid scar, we also determined additional structures of wild-type DARC (i.e. without N-terminus truncation and sortase ligation, referred to as ^WT^DARC^G42^, here onwards) in complex with CCL7 and CXCL8 at global resolutions of 3.2Å and 3.3Å, respectively (**Figure 2A-B and Supplementary Figure S1-2**). All the structures revealed a similar dimeric arrangement of the receptor with clear densities for the chemokines engaged with each receptor protomer (**Figure 2A-B and Supplementary Figure S1-2**). The transmembrane regions of the receptor exhibited sufficient resolution to model the side chains, while the proximal N-terminus (i.e. Gly^2^-Gly^39^ in CXCL8-^WT^DARC^G42^ and Gly^2^-Leu^45^ in CCL7-^WT^DARC^G42^) was not sufficiently resolved in the wild-type structures likely due to inherent flexibility. On the other hand, the longer N-terminus containing sulfated tyrosine residues was clearly resolved in the case of CXCL8-bound ^Sulfo^DARC ^G/D42^ but not in the CCL7-^Sulfo^DARC^G42^ complex. Finally, we did not observe a clear density for the seven-residue insertion resulting from sortase ligation, and therefore, this stretch was not built into the final maps of the corresponding structures. Interestingly, we observed that CCL7 engages with DARC as a monomer while CXCL8 as a dimer with only one protomer directly interacting with the receptor (**Figure 2 and Supplementary Figure S3-4**). We also observed that CXCL8-DARC complex exhibits a concatemeric assembly where the receptor protomers are linked via the free protomer of CXCL8 forming chains with multiple inter-linked dimers (**Supplementary Figure S1A-C**). However, we used only a dimeric unit for structure determination as the physiological relevance of this inter-linked multimeric assembly, if any, is not apparent currently. The overall structures of the receptor were mostly similar in these complexes with an rmsd of <1Å, the key differences were observed at the ligand-receptor interaction interface as discussed in the following sections.

**Figure 2.**
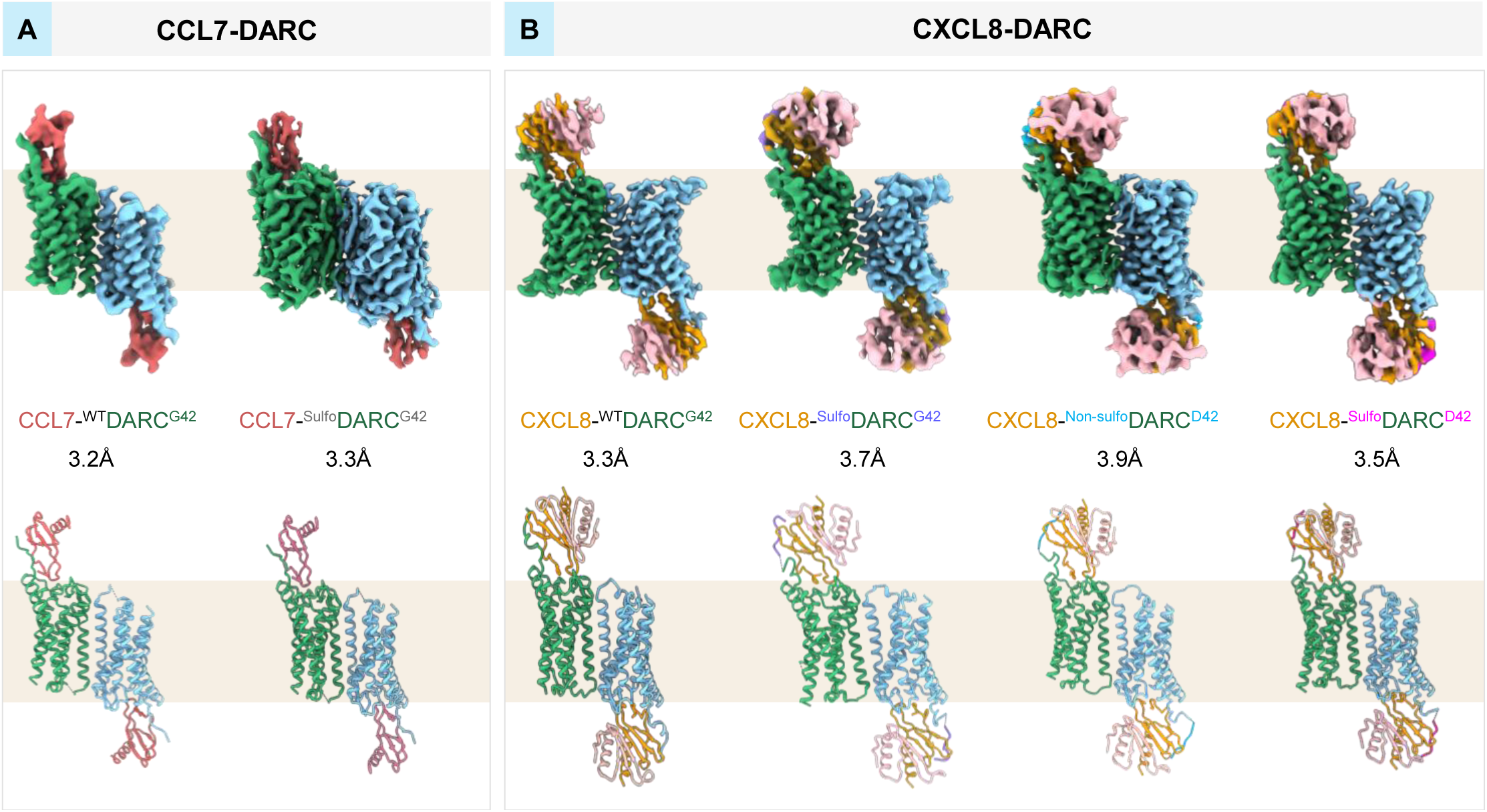
Structural snapshots of CCL7/CXCL8-DARC complexes. **(A)** Cryo-EM maps and atomic models of ^WT^DARC^G42^ and ^Sulfo^DARC^G42^ in complex with CCL7. **(B)** Cryo-EM maps and atomic models of ^WT^DARC^G42^, ^Sulfo^DARC^G42^, ^Non-sulfo^DARC^D42^ and ^Sulfo^DARC^D42^ in complex with CXCL8. The ^WT^DARC^G42^ indicates full-length, wild-type DARC without N-terminus truncation and peptide ligation.

### Molecular basis of promiscuous chemokine-binding by DARC

As mentioned earlier, we have previously reported a cryo-EM structure of CCL7-bound DARC^9^. However, the structure of CCL7-^WT^DARC^G42^ presented here exhibits a higher resolution, and therefore, has been used for a direct comparison with the CXCL8-^WT^DARC^G42^ structure. Similar to CCL7, CXCL8 also engages with ^WT^DARC^G42^ in a superficial manner reminiscent of the one-site binding mode, in contrast with the two-site binding modality observed for other chemokines with prototypical chemokine receptors (**Figure 3A**). The core of CXCL8 engages primarily with the N-terminus of ^WT^DARC^G42^ while the N-terminus of CXCL8 engages with the extracellular ends of the TM helices and extracellular loops. Interestingly, the proximal N-terminal residues of CXCL8 form a hook-like conformation, which diverges away from the orthosteric binding pocket of ^WT^DARC^G42^ (**Figure 3B**). This, in turn, precludes any significant penetration and interaction in the orthosteric binding pocket, leading to an absence of the second site of interaction.

**Figure 3.**
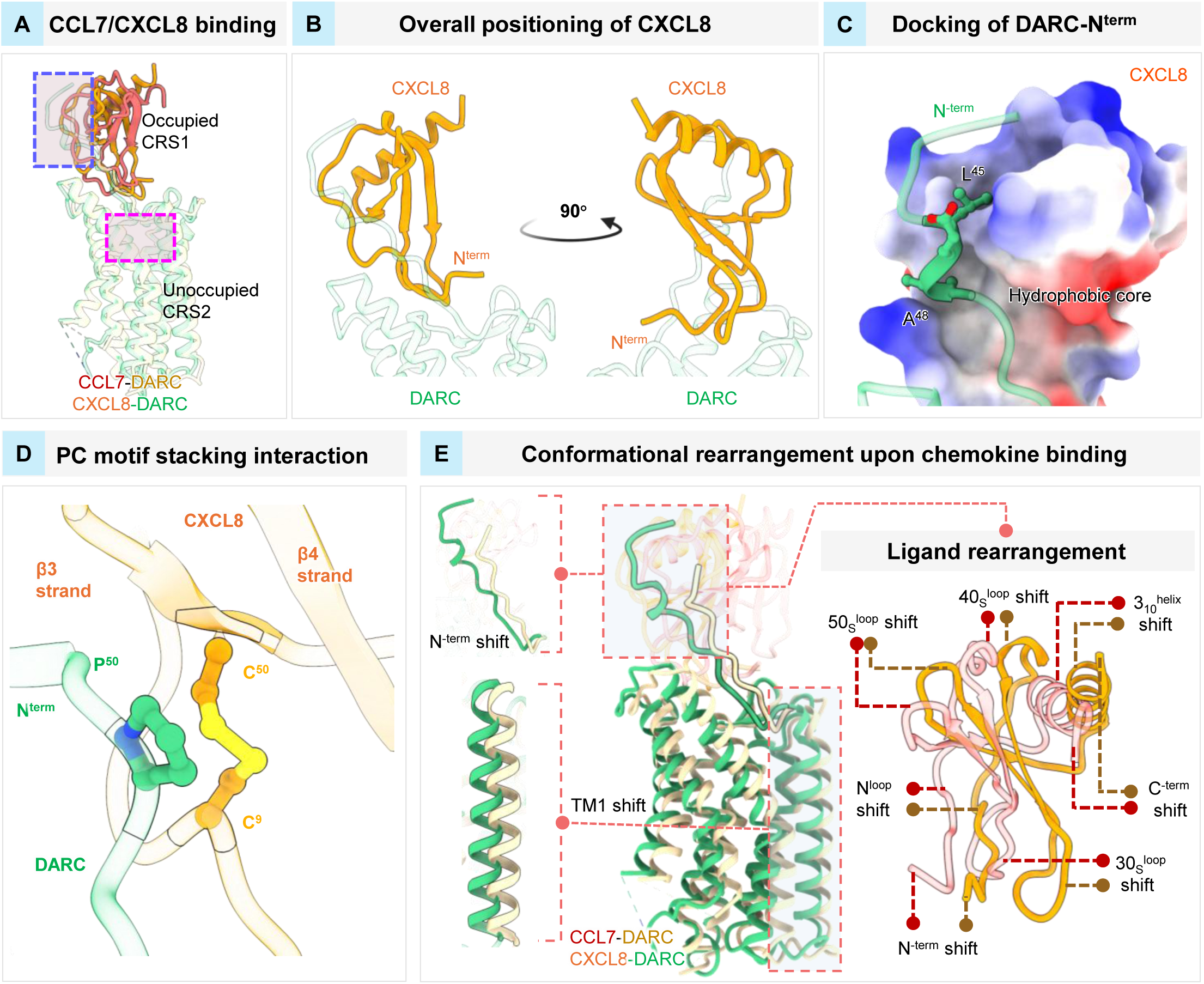
Structural differences between CCL7 vs. CXCL8 interaction with DARC. **(A)** Structural alignment of DARC-CCL7 and DARC-CXCL8 reveals a shallow binding primarily through the N-terminus of the receptor. **(B)** Conformational bending of the N-terminus of CXCL8 positions it away from the orthosteric pocket. **(C)** Docking of a short helical N-terminal segment of DARC into a hydrophobic groove of CXCL8 core domain. **(D)** Cartoon representation depicting the arrangement of P-C motif and stacking of Pro onto the Cys^9^-Cys^50^ disulfide bridge of CXCL8 **(E)** Conformational differences between CCL7 vs. CXCL8 bound DARC at the level of chemokines and the receptor. The N-terminus and TM1 of DARC shift structurally to accommodate distinct ligands (left) and the two ligands are positioned differently on DARC (right).

A larger stretch of the N-terminus of ^WT^DARC^G42^ is resolved in the CXCL8-bound structure compared to CCL7-^WT^DARC^G42^ complex, potentially suggesting a more extensive binding interface utilized by CXCL8 (**Figure 3A and Supplementary Table S2**). In particular, Ala49^N-term^-Asn55^N-term^ segment of ^WT^DARC^G42^ forms extensive interactions with the N^loop^ of CXCL8, whereas the Leu45^N-term^-Ala48^N-term^ segment adopts a helical conformation and docks into a hydrophobic pocket lined by β3-strand residues in the CXCL8 core domain (**Figure 3C and Supplementary Figure S5A**). We observe that a network of hydrogen bonds, hydrophobic contacts, and ionic interactions stabilize the interaction between CXCL8 and the N-terminus of ^WT^DARC^G42^. For example, Asp46^N-term^ and Ser53^N-term^ form hydrogen bonds with Arg47^CXCL8^ and Arg6^CXCL8^, respectively, while Tyr41^N-term^ engages Lys15^CXCL8^ through a polar contact. Moreover, hydrophobic interactions link Leu45^N-term^ to Leu43^CXCL8^ and Leu49^CXCL8^, and Asp40^N-term^ makes a non-bonded contact with Leu43^CXCL8^ **(Supplementary Table S2)**.

The P-C motif is a conserved Pro-Cys cluster present in the N-terminus of certain chemokine receptors that enhances ligand subgroup selectivity and helps induce a kink in the N-terminus, facilitating proper folding and orientation against the chemokines^9,41^. Pro50^N-term^ of the P-C motif in ^WT^DARC^G42^ stacks against the Cys9^CXCL8^-Cys50^CXCL8^, thus forming one of the critical interaction sites at CRS1 (Chemokine Recognition Site 1) (**Figure 3D**). In addition, the binding includes hydrogen bonds between Glu4^CXCL8^ and Leu5^CXCL8^ of the CXCL8 N^loop^ with Ser53^N-term^ and Arg267^6.62^, respectively. Additionally, Pro32^CXCL8^, His33^CXCL8^, and Asn36^CXCL8^ of the CXCL8 30s^loop^ make non-bonded contacts with Tyr199^ECL2^, Arg267^6.62^, and Lys269^6.64^ respectively. The N-terminus of ^WT^DARC^G42^ undergoes a linear transition to engage with the core domain of CXCL8 compared to CCL7 (**Supplementary Table S2**). The 40s^loop^ of CXCL8 protomer interacting with ^WT^DARC^G42^ rotates and translates by ∼35° and ∼8Å, respectively, relative to CCL7, which in turn repositions the N^loop^ of CXCL8 on the extracellular surface of the receptor (**Figure 3E**). The N-terminus of CCL7 is positioned at a shallower depth in the orthosteric pocket than CXCL8 (∼21Å vs. ∼17Å from Trp^6.48^), and even more so compared to that observed for CXCL8 in complex with a prototypical chemokine receptor, CXCR2 (∼13Å) (**Supplementary Figure S5B**).

In order to validate the contribution of key residues in ^WT^DARC^G42^ implicated in differential recognition of CCL7 and CXCL8, we performed alanine substitutions at Glu46^N-term^, Ser53^N-term^, Arg267^6.62^, Ser274^ECL3^, and Asp283^7.32^ based on structural analyses of CCL7- and CXCL8-bound ^WT^DARC^G42^ structures (**Figure 4A-B**). We observed that Arg^267^Ala and Asp^283^Ala substitutions reduced CXCL8 binding without affecting CCL7 interaction with ^WT^DARC^G42^, which aligns with the structural observations. Arg267^6.62^ in ^WT^DARC^G42^ forms a hydrogen bond with Leu^5^ in CXCL8, whereas it engages only in non-bonded contacts with Ser^34^ and His^35^ in CCL7. Similarly, Asp283^7.32^ in ^WT^DARC^G42^ does not interact with CCL7 but it forms a hydrogen bond and a salt bridge with Arg6 in CXCL8. In contrast, Glu^46^Ala and Ser^53^Ala mutations selectively impaired CCL7 binding with ^WT^DARC^G42^ without affecting the interaction with CXCL8. Glu46 in the N-terminus of ^WT^DARC^G42^ forms a salt bridge with Lys^18^ in CCL7, however, this interaction is absent in the CXCL8-^WT^DARC^G42^ complex. Similarly, Ser^53^ in the N-terminus of ^WT^DARC^G42^ interacts primarily with Ser^8^ in CCL7, and therefore, its mutation impairs CCL7 binding to ^WT^DARC^G42^. However, in case of CXCL8-^WT^DARC^G42^, despite Ser^53^ contacting the main chain of Arg^6^ of CXCL8, additional contacts with Cys^51^, Gln^7.28^ and Asp^7.32^ of ^WT^DARC^G42^, further stabilise this interaction, and therefore, alanine mutation of Ser53 does not dramatically alter CXCL8 binding. Finally, mutation of Ser^274^ in ^WT^DARC^G42^ does not appear to impact the binding of either of the chemokines as it primarily engages Lys^11^ and Tyr^13^ in CXCL8 and CCL7, respectively, vias main-chain contacts (**Figure 4A-B**). Taken together, these observations suggest that the conformational plasticity of the N-terminus of DARC may allow a tunable positioning of different chemokines on the receptor, thereby driving promiscuous chemokine engagement. This is further supported by the observation that the N-terminus of DARC is the longest among all the chemokine receptors, perhaps designed to accommodate different chemokines in a modular fashion (**Figure 4C**). It is also interesting to note that the orthosteric binding pocket in DARC exhibits maximal deviation in terms of the nature of amino acid residues compared to the other chemokine receptors, which likely provides an incompatibility for engaging the N-terminus of chemokines, thereby, imparting one-site binding modality unlike other chemokine-chemokine receptor pairs (**Figure 4D**).

**Figure 4.**
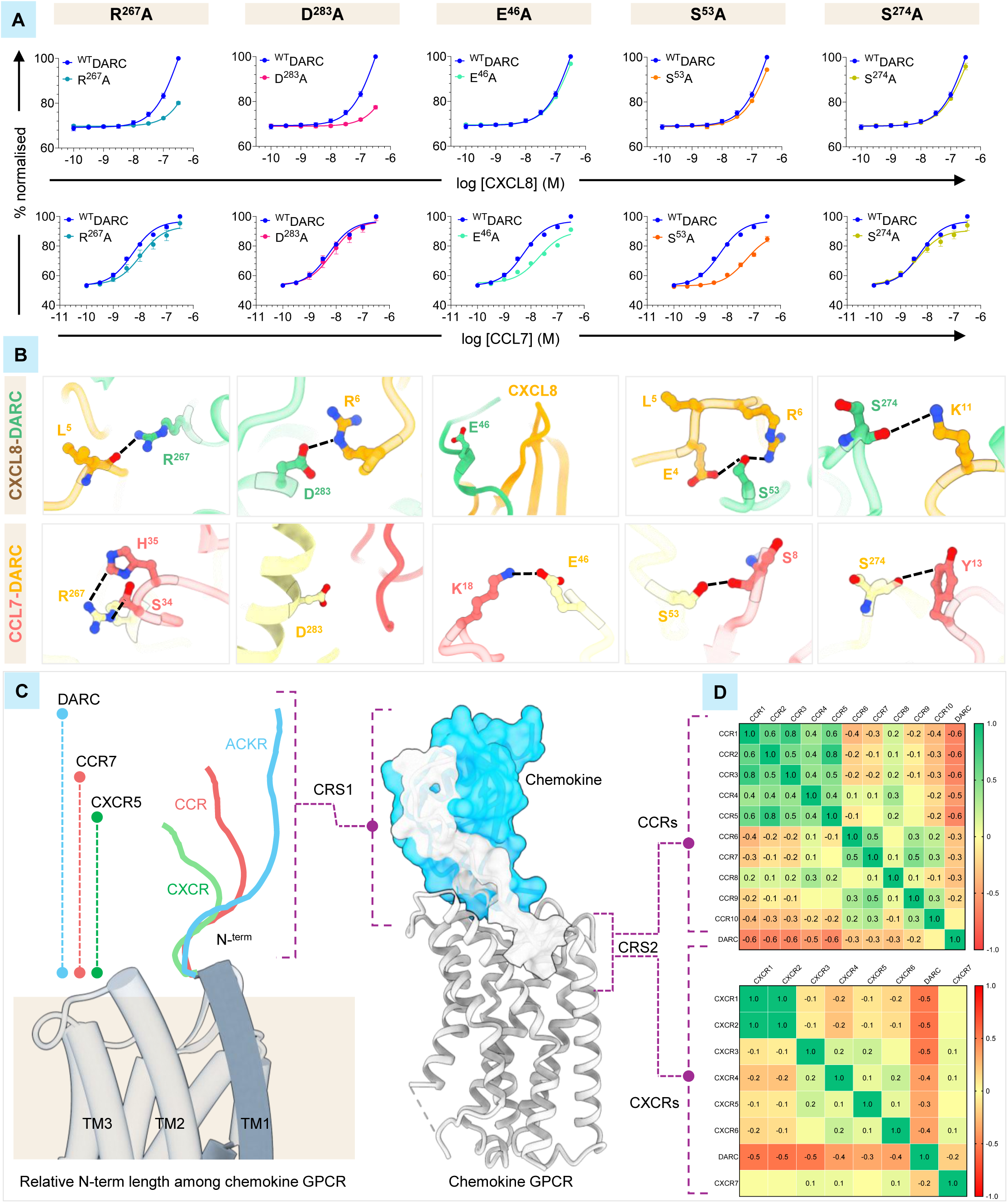
Structural insights into promiscuous chemokine interaction with DARC. **(A)** Effect of alanine substitutions at the indicated residues in DARC on CXCL8 and CCL7 binding assessed using a NanoBRET-based binding assay. **(B)** Structural correlation of CXCL8 and CCL7 binding to indicated mutants of DARC visualized based on the structural snapshots. **(C)** A schematic representation of N-terminus length across all chemokine receptors to highlight the longest N-terminus of DARC. **(D)** Sequence variation in CRS2 (chemokine recognition site 2) across the C-C and C-X-C chemokine receptors including DARC. The color coding (red to green) indicates most to least sequence divergence.

### Atypical structural features of DARC

As CXCL8 is a common ligand for DARC and CXCR2, we next compared the structures of CXCL8-bound CXCR2 (PDB: 8XWN)^27^ and ^WT^DARC^G42^. The structural superimposition indicates an overall similar receptor structure with an rmsd of <1.2Å across the main-chain (**Figure 5A**), still however, there are notable differences. For example, in the CXCL8-DARC complex, the N-terminal stretch spanning Leu^45^-Ala^49^ exhibits a lateral shift of ∼12Å compared to CXCR2 (**Supplementary Figure S5C**) while the extracellular end of TM1 and the distal N-terminus of ^WT^DARC^G42^ are positioned towards the orthosteric binding pocket compared to CXCR2. In addition, in the CXCL8-^WT^DARC^G42^ structure, the extracellular end of TM2 shortens, ECL1 shifts outward by ∼7Å, and ECL2 is also displaced outward from the orthosteric site by ∼12 Å compared to that in CXCL8-CXCR2 (**Supplementary Figure S5C**). These changes culminate into a narrower orthosteric binding pocket in compared to CXCR2 (i.e. buried surface area of ∼4000Å^2^ vs. ∼5000 Å^2^, respectively), and in turn, impose a steric hindrance limiting the penetration of CXCL8 in the orthosteric binding pocket (**Supplementary Figure S6A**). Such differences probably underlie distinct chemokine N-terminal binding poses between DARC and CXCR2.

**Figure 5.**
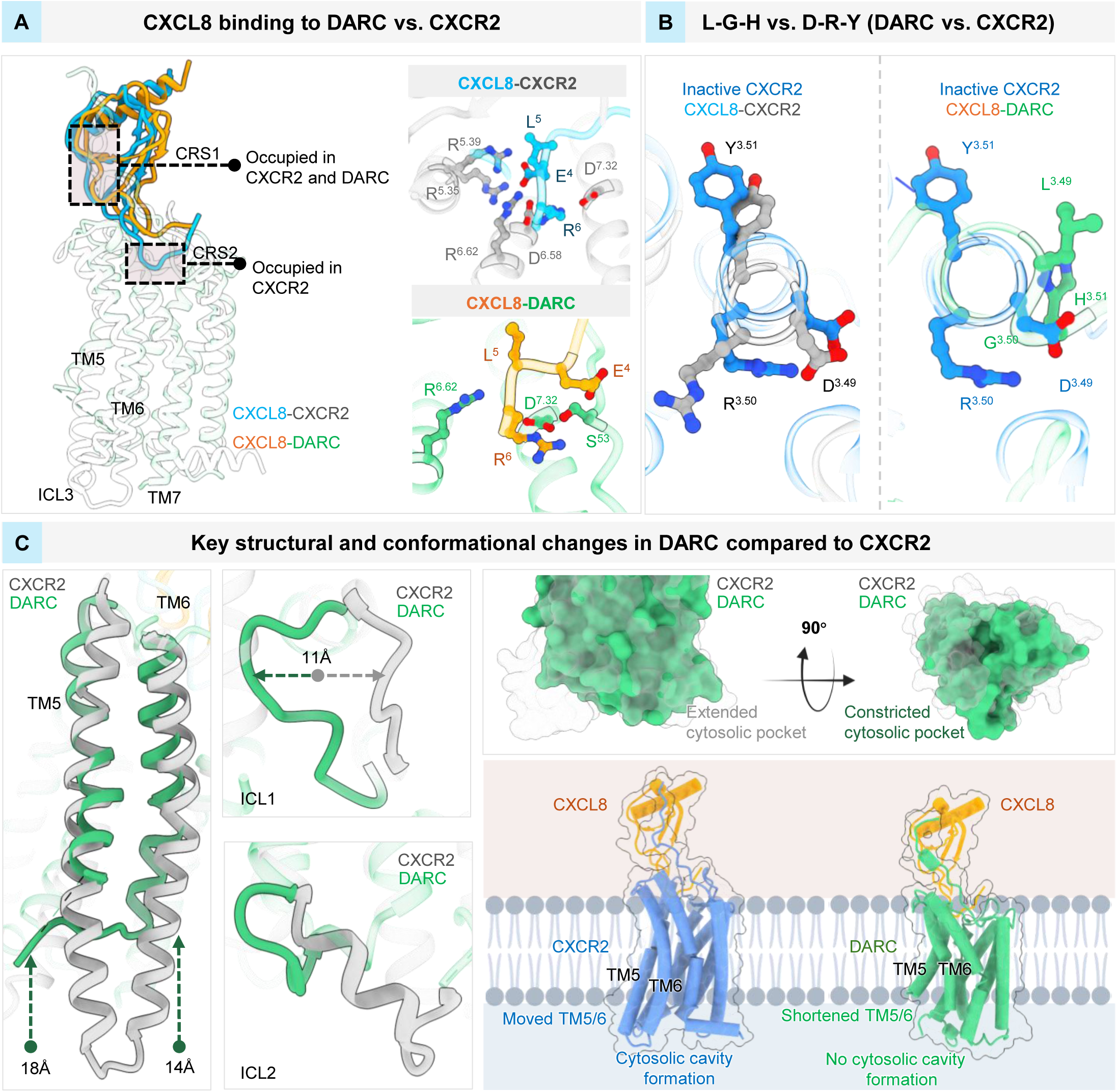
Structural comparison of CXCR2 and DARC. **(A)** Superimposition of CXCL8-bound DARC and CXCR2 structures elucidate a relatively superficial binding mode on DARC compared to CXCR2 (left panel). The E-L-R-motif in CXCL8 engages with a partial R-D motif in DARC unlike a full R-D motif in CXCR2. **(B)** DARC harbors an L-G-H sequence instead of the conserved D-R-Y motif in CXCR2 with distinct structural positioning as visualized in the structural snapshots. **(C)** Shortening of TM5 and TM6, together with conformational shifts in ICL1 and ICL2, likely preclude the formation of a defined cytoplasmic pocket to engage canonical signal-transducers.

We have recently identified a specific interaction between an E-L-R sequence motif in C-X-C chemokines with a structural R-D motif in CXCR2 as a molecular mechanism driving promiscuous chemokine binding^27,28^. That is, the linear E-L-R motif in CXCL8 (i.e. Glu^4^-Leu^5^-Arg^6^) engages with a structural R-D motif organized in CXCR2 involving Arg208^5.35^, Arg212^5.39^, Asp274^6.58^, Arg278^6.62^ and Asp293^7.32^ ^27^. Interestingly, we observe the formation of only a partial R-D motif in ^WT^DARC^G42^ due to substitution at the positions corresponding to Arg^5.35^ and Arg^5.39^ with Leu^5.35^ and Gln^5.39^, and an upward displacement of the E-L-R motif containing segment in CXCL8 along the vertical axis of the receptor (**Figure 5A**). These structural features may represent an adaptive mechanism in DARC that allows superficial binding of chemokines, and thereby, imparting promiscuous recognition.

Finally, DARC lacks micro-switches that are typically conserved in prototypical GPCRs including CXCR2, that facilitate receptor activation, although it retains variants of these. For example, instead of the conserved D-R-Y motif at the cytoplasmic end of TM3, which forms an ionic-lock with TM6 in CXCR2, DARC harbors an L-G-H sequence (**Figure 5B**). Interestingly, we also observe a ∼65° kink at the cytoplasmic end of TM3 and ICL2 in CXCL8-^WT^DARC^G42^ compared to CXCL8-CXCR2 which precludes the formation of ionic-lock with TM6 (**Figure 5B**). Similarly, DARC also harbors T-P-x-x-L in TM7 instead of N-P-x-x-Y, and C-F-x-P in TM6 instead of C-W-x-P, which impart distinct intra-helical arrangements in DARC compared to CXCR2 and other GPCRs (**Supplementary Figure S6B)**. Furthermore, the P-I-F motif of CXCR2 and other GPCRs, implicated in relaying the signal from the extracellular to intracellular by allosterically linking ligand-binding to G protein-coupling, is also altered in DARC displaying instead an A-F-W sequence (**Supplementary Figure S6B)**. Additionally, TM5 in CXCL8-^WT^DARC^G42^ exhibits a shortening by ∼18Å, whereas TM6 exhibit a shortening by ∼14Å, compared to CXCL8-CXCR2 structure (**Figure 5C**). Finally, there are distinct conformational changes on the intracellular side of DARC compared to CXCR2. For example, the ICL1 in DARC shifts laterally by ∼11Å, which will likely displace H8, and ICL2 relocates due to shortened TM3, and TM5 and TM6 (**Figure 5C**). Collectively, these structural changes nearly abolish the formation of the cytoplasmic pocket on DARC, rationalizing the lack of engagement with canonical signal-transducers such as G-protein, GRKs, and βarrs. We note that these observations are similar to those in the CCL7-bound structure reported previously^9^, and also evident in the higher resolution CCL7-^WT^DARC^G42^ structure presented here.

### Modulation of chemokine binding by Fy^a^ vs. Fy^b^ allelic variation and tyrosine sulfation

Fy^a^ and Fy^b^ allelic variants differ at the level of single amino acid residue at position 42 (i.e. Gly^42^Asp), which may alter the electrostatic properties at the interface of chemokine binding to DARC. In fact, previous studies have demonstrated that DARC^D42^ (Fy^b^) variant has a greater sequestration capacity for pro-inflammatory chemokines such as CCL2, CCL5 and CXCL8 compared to DARC^G42^ (Fy^a^), implicating allele-specific binding differences that might alter local chemokine gradients and inflammatory responses^29,42^. A closer analysis of the cryo-EM structural snapshots reveals that in the Fy^a^ variant of DARC, the N-terminal segment retains greater conformational plasticity around Gly^42^, allowing the N-terminal loop to adopt a more relaxed orientation relative to the CXCL8 core domain. In contrast, the Fy^b^ exhibits a more rigid N-terminal hinge due to Asp^42^ substitution, which also induces a lateral displacement and repositioning of this segment of DARC towards CXCL8 core domain by ∼3.3 Å (**Figure 6A**). This repositioning of DARC N-terminus in Fy^b^ variant is accompanied by substantial reorientation of CXCL8 interacting residues, especially, Asp^42^, which forms an ionic interaction with Lys^15^ in CXCL8 (**Figure 6A**).

**Figure 6.**
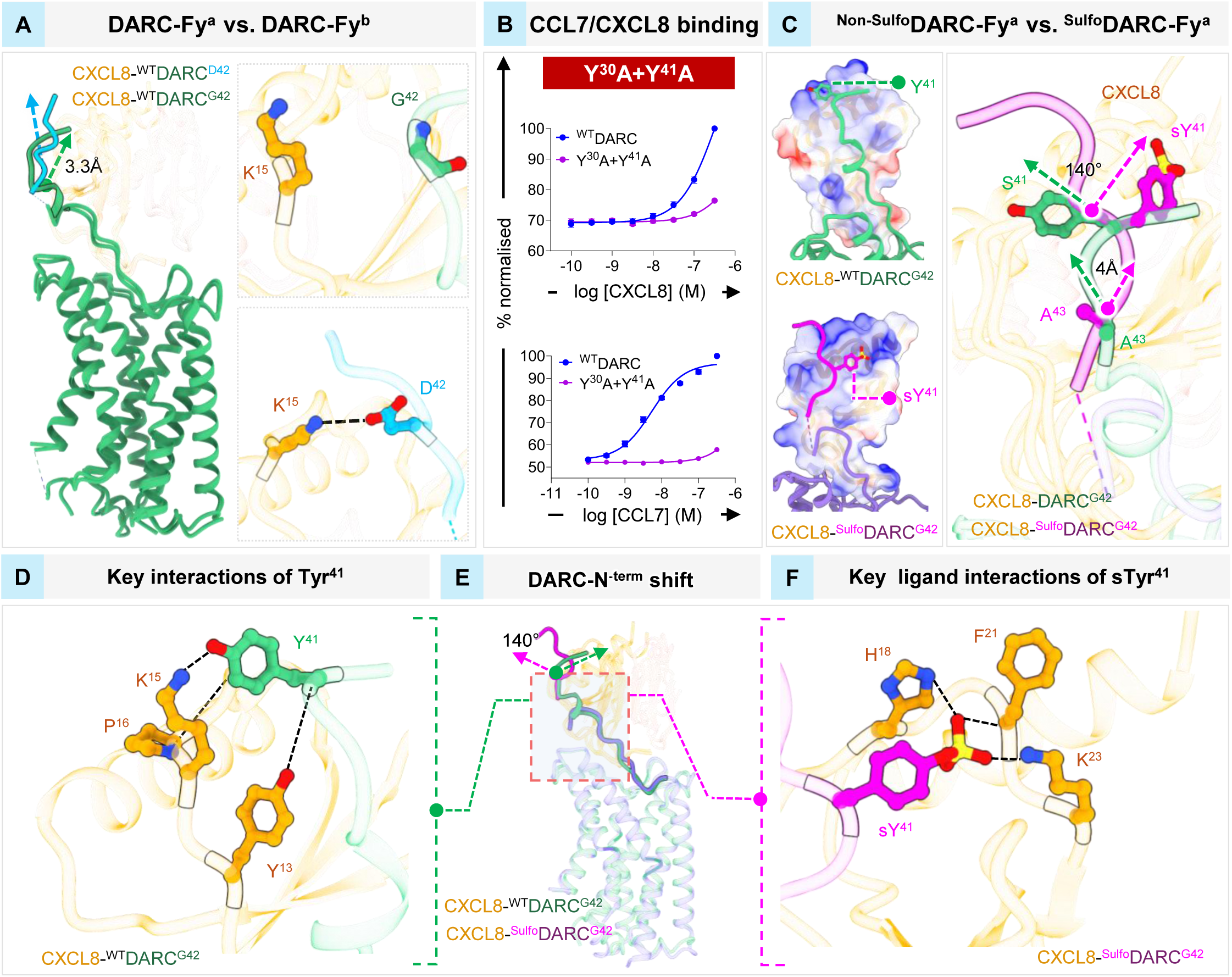
Effect of tyrosine sulfation on chemokine-binding to DARC. **(A)** Conformational rearrangements of DARC due to Fy^a^ vs. Fy^b^ polymorphism. **(B)** Effect of Tyr^30+41^Ala substitution on CCL7 and CXCL8 interaction as assessed by NanoBRET binding assay. **(C)** Repositioning of the N-terminus of the receptor in ^Non-Sulfo^DARC and ^Sulfo^DARC upon interaction of CXCL8. **(D-F)** Differential engagement of Tyr^41^ in ^Non-Sulfo^DARC and ^Sulfo^DARC with the key residues in CXCL8.

Previous studies have reported distinct contributions of tyrosine sulfation in the N-terminus of several chemokine receptors including DARC on chemokine binding^14,29^. The N-terminus of DARC harbors two putative tyrosine sulfation sites namely Tyr^30^ and Tyr^41^, and we first assessed the binding of CCL7 and CXCL8 with the wild-type receptor and a mutant harboring Tyr^30^Ala and Tyr^41^Ala substitutions (Tyr^30+41^Ala). We observed that the binding of both, CCL7 and CXCL8, was impaired for the Tyr^30+41^Ala mutant compared to the wild-type receptor (**Figure 6B**). Next, we examined the cryo-EM structures of ^Sulfo^DARC^G42^ in complex with CXCL8, however, as mentioned earlier, we observed discernible density of only sulfated Tyr^41^ in the CXCL8-^Sulfo^DARC^G42^ complex while the stretch harboring sulfated Tyr^30^ was not resolved at sufficient level to model it in the structure. The overall binding and positioning of CXCL8 on non-sulfated and sulfated DARC are similar, however, there are critical differences around Tyr^41^ residue in terms of the interaction network **(Supplementary Table S4-S5)**. As discussed earlier, in the CXCL8-^WT^DARC^G42^ structure, the Leu^45N-term^-Ala^48N-term^ segment adopts a short helical conformation, which docks into a hydrophobic pocket lined by β3-strand residues of the core domain of CXCL8. On the other hand, in the CXCL8-^Sulfo^DARC^G42^ structure, the N-terminal residues preceding Ala^43N-term^ deviate by ∼4Å from the core of CXCL8 (**Figure 6C**). As a result, the side-chain of sulfated Tyr^41^ rotates by ∼140° towards the CXCL8 core domain compared to the side-chain of non-sulfated Tyr^41^ (**Figure 6D-F**). Concomitantly, Phe^21^ in the N-loop of CXCL8 moves away from the core domain by ∼80° to create a sub-pocket that accommodates sulfated Tyr^41^. Unlike non-sulfated Tyr^41^, the sulfated Tyr^41^ engages with His^18^ and Lys^23^ in CXCL8 via direct ionic-interactions stabilizing its position in the sub-pocket (**Figure 6D**). Moreover, Asp^40^ and Glu^46^ in the N-terminus of ^Sulfo^DARC rotate by ∼140° and ∼100°, respectively, away from the core of CXCL8, and engage with Lys^15^ and Lys^11^ in the ligand **(Supplementary Figure S7A**).

Comparison of the Fy^b^ variant of DARC in the presence and absence of tyrosine sulfation reveals additional fine-tuning of chemokine interactions. In the CXCL8-bound non-sulfated Fy^b^ form, Tyr^41^ engages Leu^43^, Asp^45^ and Arg^47^ of CXCL8, however, upon sulfation, sTyr^41^ reorients to establish interactions with His^18^, Phe^21^, Lys^23^ and Ser^44^ of CXCL8 (**Figure 7A**). Simultaneously, Asp^42^ within the N-terminal segment rotates towards the CXCL8 core by approximately ∼110° upon Ty41 sulfation (**Figure 7A**). Moreover, the N-terminal segment preceding sTyr^41^ undergoes a lateral displacement of ∼5.6 Å towards the CXCL8 core domain (**Figure 7A**). Collectively, these changes indicate that tyrosine sulfation does not merely strengthen binding through additional electrostatic contacts but actively remodels the interaction interface between the Fy^b^ variant and CXCL8. The structural comparison of CXCL8-bound Fy^a^ vs. Fy^b^ variants of ^Sulfo^DARC reveals that allelic substitutions further reshape the conformational dynamics of the N-terminus of DARC while preserving the overall mode of chemokine recognition. For example, the N-terminal segment preceding sTyr^41^ exhibits a substantial conformational repositioning in Fy^b^ compared to Fy^a^, reflected by a shift of ∼9 Å of this segment toward the core domain of CXCL8. As a result, Asp^38^ in the Fy^b^ variant extends linearly towards the CXCL8 core domain and forms an interaction with Lys^20^ of CXCL8, an interaction that is absent in Fy^a^ **(Supplementary Figure S7B**). In contrast, the N-terminal segment of the receptor following of sTyr^41^ exhibits increased rigidity in Fy^b^, as Asp^42^ stabilizes the hinge region and allows direct interactions with Tyr^13^ and Arg^47^ of CXCL8 (**Figure 7B**). On the other hand, in Fy^a^, Asn^44^ compensates for these missing interactions to some extent by engaging Tyr^13^ of CXCL8 and properly aligning the N-terminus of CXCL8 along the receptor N-terminus (**Figure 7B**).

**Figure 7.**
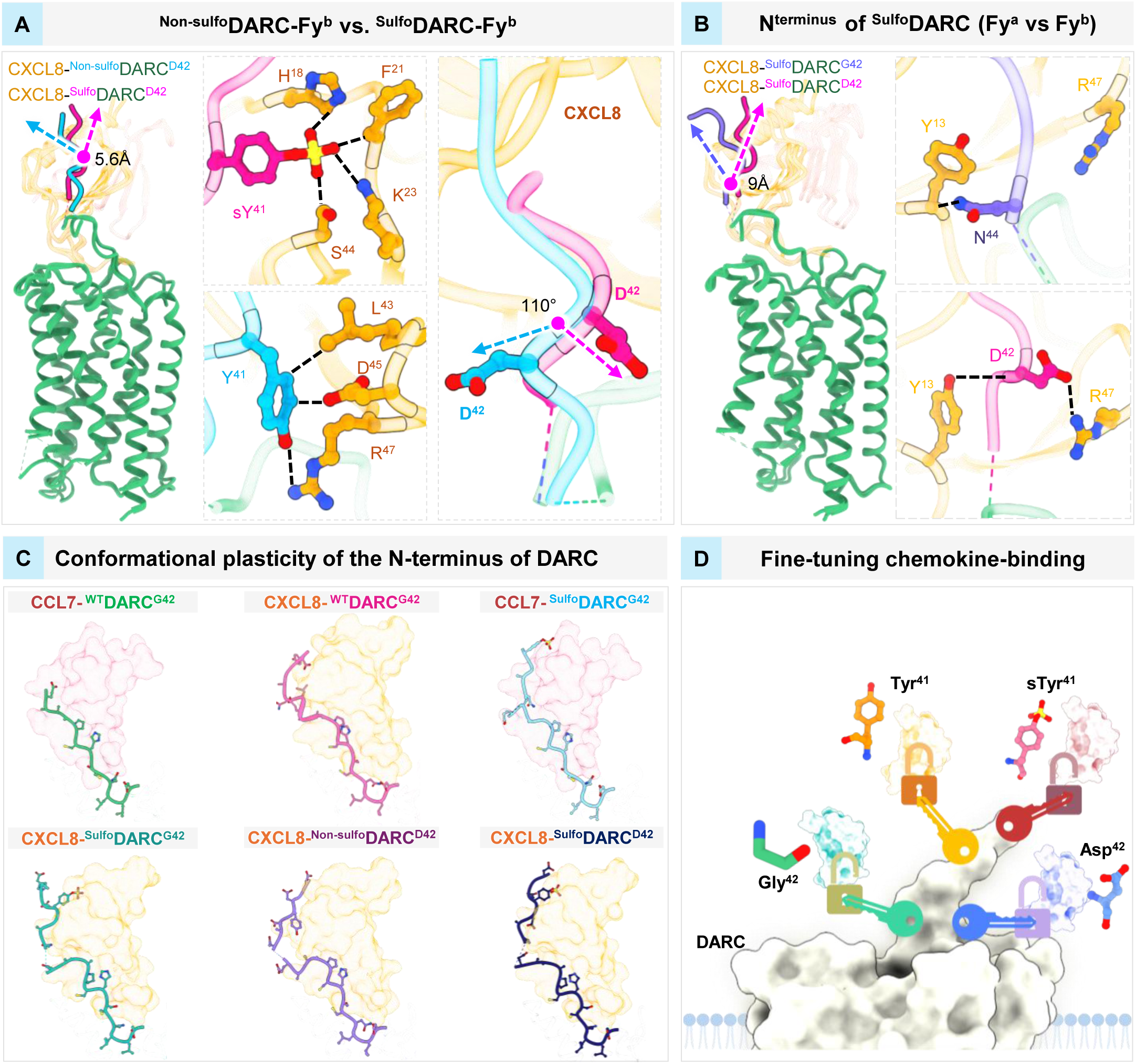
Interplay of tyrosine sulfation and Gly^42^Asp polymorphism on chemokine binding. **(A)** Differential interaction network of sTyr^41^ vs. Tyr^41^ in the Fy^b^ variant of DARC with CXCL8. **(B)** Key differences in the positioning and interaction of the N-terminus of DARC between the sulfated versions of Fy^a^ vs. Fy^b^ variants. (**C)** Cartoon representation showing conformational plasticity of the N-terminus of DARC in complex with different chemokines, upon tyrosine sulfation, and Fy^a^/Fy^b^ allelic variations**. (D)** A schematic representation depicting the key determinants in the N-terminus of DARC, which fine-tune promiscuous chemokine-engagement using a conserved binding modality.

## Discussion

We have previously demonstrated a near-complete absence of canonical transducer-coupling to DARC including G-protein activation, GRK and β-arrestin recruitment, and second messenger response^9^. Based on a cryo-EM structure of CCL7-DARC, we attributed this functional divergence primarily to the lack of effector-site engagement by CCL7, and shortened TM5-TM6 topology incompatible with stable interaction of canonical transducers^9^. Interestingly, the structural snapshots of DARC in complex with CXCL8 presented here further corroborate the same observation, suggesting this as the intrinsic mechanism irrespective of interacting chemokine. The primary engagement of chemokines with the N-terminus of DARC without a significant interaction in the orthosteric binding pocket offers plausible explanation for promiscuous chemokine binding. For example, the conformational plasticity of the N-terminus of DARC observed here in various structures reveals that an on-demand adjustment and positioning of the chemokines may occur on the receptor, which is further fine-tuned by tyrosine sulfation to stabilize their interactions. This, taken together with no strict dependence on orthosteric binding site engagement, likely imparts complete flexibility to accommodate different chemokines considering an overall similar domain architecture. This is further substantiated by the observation that the nature of amino acids in the orthosteric binding pocket of DARC is most divergent among the chemokine receptors, and the interaction of CXCL8 with CXCR2 involving a direct engagement of the E-L-R motif in chemokines and the R-D motif in the receptor is also not conserved in DARC. This molecular mechanism yielding promiscuous chemokine binding may underlie the ability of DARC to serve as a chemokine scavenger for a broad repertoire of chemokines, and also transcytosis functions proposed previously^27^.

Although DARC exhibits promiscuous binding to several chemokines, they are likely to differ in their binding affinity for the receptor^5,19,43,44^. It is possible that despite binding primarily through the N-terminus, different chemokines have distinct sensitivity for tyrosine sulfation on the receptor^31–36^. This hypothesis is supported by our observations that sulfo-Tyr^41^ is not resolved as well in CCL7-^Sulfo^DARC complex as in the CXCL8-^Sulfo^DARC structure and is further corroborated by site-directed mutagenesis data. In addition to chemokines, the interaction of the Duffy binding protein of *P. vivax* (PvDBP) with DARC also requires N-terminal sulfation^10–14^, and therefore, the chemical-ligation strategy presented here provides a tool for capturing and visualizing the PvDBP-DARC complex in future studies. We also note that the chemical-ligation strategy utilized here leaves a minor seven amino acid scar on the receptor. While this does not appear to influence the binding modality of chemokines, future development of tailored chemical-ligation strategies, for example, may overcome this limitation.

In conclusion, the work described in this manuscript has revealed key molecular insights into promiscuous chemokine recognition by DARC and modulation by N-terminal tyrosine sulfation (**Fig. 7C-D**). These findings should facilitate a better understanding of distinct functional divergence displayed by DARC and provide a framework that could potentially be leveraged to design novel therapeutics.

## Acknowledgements

The research on Duffy antigen receptor for chemokines (DARC) in our laboratories is supported by Department of Biotechnology (BT/PR47867/COT/142/59/202), the Australian Research Council (Laureate Fellowship FL250100011 to RJP), and JSPS KAKENHI grant numbers JP23KJ0491 (F.K.S.); the Japan Agency for Medical Research and Development (AMED) grant numbers JP25gm7110004 (F.K.S.) and JP223fa627001 (F.K.S.). A part of the cryo-EM data was collected at the National cryo-EM Facility at IIT Kanpur established with the support from ANRF/SERB (IPA/2020/000405). In addition, the research program on chemokine receptors in our laboratory is also supported by the Senior Fellowship of the DBT Wellcome Trust India Alliance (IA/S/20/1/504916), the Indian Council of Medical research (EMDR/CAR/13/2024-01-00013), and the Department of Science and Technology (DST/TTI/TC/AMR/COE/2023/5), J.C. Bose Grant from ANRF (ANRF/JBG/2025/000055/LS), (BT/PR53019/MED/30/2528/20) and LADY TATA Memorial Trust. A.K.S. is the Sonu Agrawal Memorial Chair Professor. This study was supported in part by the Luxembourg Institute of Health (LIH) through the NanoLux Platform to AC. We sincerely thank Prof. Volodymyr Korkhov, and Drs. Basavraj Khanppnavar, Jagannath Maharana, and Shirsha Saha for their contribution to the early efforts on CXCL8-DARC structure determination.

## Authors’ contribution

MG, NR, DT expressed and purified the receptor and reconstituted the complexes for structural analysis; SK synthesized the sulfated peptides for ligation with guidance from RJP and MS; AD, MKY, SM, and NB contributed to expression and purification of the receptor and ligands; YM, KS, KMH, KA collected and processed cryo-EM data on CCL7-^WT^DARC^G42^, CXCL8-^WT^DARC^G42^ and CCL7-^Sulfo^DARC^G42^ complexes and determined the structures with overall guidance from FKS; AC carried out the mutagenesis and chemokine binding experiments; MG and RB processed the cryo-EM data of the CXCL8-^Sulfo^DARC^G42^, CXCL8-^Sulfo^DARC^D42^, CXCL8-^Non-sulfo^DARC^D42^ complexes, determined the structures, carried out the structural analysis and prepared the figures; all authors contributed to writing and editing the manuscript; AC, RJP, ON and AKS supervised the overall project.

## Data availability

The cryo-EM maps and structures have been deposited in the EMDB and PDB with accession numbers EMD-67876, PDB ID-21OZ (CXCL8-^WT^DARC^G42^); EMD-67880, PDB ID-21PE (CXCL8-^Sulfo^DARC^G42^-composite map); EMD-67890 (CXCL8-^Sulfo^DARC^G42^-receptor focused map); EMD-67889 (CXCL8-^Sulfo^DARC^G42^-ligand focused map); EMD-67879, PDB ID-21PD (CXCL8-^Sulfo^DARC^D42^-composite map); EMD-67885 (CXCL8-^Sulfo^DARC^D42^-full map); EMD-67887 (CXCL8-^Sulfo^DARC^D42^-ligand focused map); EMD-67877, PDB ID-21PB (CXCL8-^Non-sulfo^DARC^D42^-composite map); EMD-67882 (CXCL8-^Non-sulfo^DARC^D42^-full map); EMD-67883 (CXCL8-^Non-sulfo^DARC^D42^-ligand focused map); EMD-67874, PDB ID-21OX (CCL7-^WT^DARC^G42^); and EMD-67873, PDB ID-21OW (CCL7-^Sulfo^DARC^G42^).

## Materials and methods

### General chemicals and reagents

Most of the general reagents were purchased from Sigma-Aldrich unless otherwise specified. Dulbecco’s Modified Eagle’s Medium (DMEM), Trypsin-EDTA, Fetal Bovine Serum (FBS), Phosphate-Buffered Saline (PBS), Hank’s Balanced Salt Solution (HBSS), and Penicillin-Streptomycin solution were obtained from Thermo Fisher Scientific. HEK-293T cells (ATCC) were maintained in DMEM (Gibco, Cat. No: 12800-017) supplemented with 10% (v/v) FBS (Gibco, Cat. No: 10270-106) and 100 U/mL penicillin and 100 μg/mL streptomycin (Gibco, Cat. No: 15140122) at 37 °C under 5% CO₂. *Sf9* cells were cultured in protein-free media (Gibco, Cat. No: 10902-088) at 27 °C at 135 rpm.

### Human cell line

HEK-293T cells were purchased from Abcam and grown in Dulbecco’s modified Eagle medium (DMEM) supplemented with 10% fetal bovine serum (Sigma) and penicillin/streptomycin (100 Units per ml and 100 μg per ml). HEK293T cells stably expressing untagged DARC or DARC N-terminally fused to Nanoluciferase were established using pIRES-puromycin vectors and were grown in DMEM medium supplemented with 10% fetal bovine serum (Sigma) and penicillin/streptomycin (100 Units per ml and 100 μg per ml) and puromycin (5 μg per ml).

### Insect cell line

*Spodoptera frugiperda (Sf9)* cells were obtained from Expression systems. The cells were maintained in glass conical flasks at a density of 0.9 million cells per mL in protein-free insect cell medium with regular splitting at every alternate day. These cells were grown in a shaker incubator at 27 °C with a constant agitation at 135 rpm.

### Bacterial cell culture

*Escherichia coli* strain DH5-alpha were used for plasmid DNA amplification and isolation, and they were cultured in Luria-Bertani (LB) broth at 37 °C with shaking at 160 rpm. For protein expression, BL21 (DE3), Rosetta (DE3), SHuffle strains of *Escherichia coli* were used, and they were cultured using Luria-Bertani (LB), Terrific Broth (TB), or 2XYT media under the indicated culture conditions as described in the subsequent method sections.

## Methods

### Preparation of unmodified or sulfated DARC N-terminal peptides

The peptide was prepared on a pre-loaded Fmoc-Ala Wang resin (100 µmol) using Fmoc-SPPS on a SYRO I automated peptide synthesizer. Wang resin in polypropylene reaction vessels was Fmoc-deprotected with 40% (v/v) piperidine in dimethylformamide (DMF) (800 μL) for 3 min with intermittent shaking, drained by vacuum, and treated with 20% (v/v) piperidine in DMF (800 μL) for 10 min, drained, and washed with DMF (4 × 1.2 mL). The resin was then treated with a solution of Fmoc-AA-OH (4 eq.) and Oxyma (4.4 eq.) in DMF (400 μL), followed by the addition of a 1% (w/v) solution of 1,3-diisopropyl-2-thiourea in DMF (400 μL), and a solution of *N,N′-*diisopropylcarbodiimide (4 eq.) in DMF (400 μL). Fmoc-Tyr(SO_3_nP)-OH (nP = neopentyl) was synthesized according to the literature protocol^33^ and was coupled at room temperature manually using the aforementioned DIC/Oxyma coupling conditions (with 1.5 eq. of Fmoc-Tyr(SO_3_nP) used per coupling). Coupling reactions were conducted at appropriate temperature (15 min at 75 °C for unmodified DARC peptide; 30 min at 40 °C for sulfated DARC peptide) using a heating block and intermittently shaken. The resin vessel was then vacuum drained and washed with DMF (4 × 1.2 mL). A capping step was performed with a solution of 5% (v/v) acetic anhydride (Ac_2_O) and 10% (v/v) N,N-diisopropylethylamine (*i*Pr_2_NEt) in DMF (800 μL) for 6 min with intermittent shaking at room temperature following each coupling reaction. The resin vessel was then vacuum drained and washed with DMF (4 × 1.2 mL). Iterative cycles of this process were repeated until complete peptide elongation was achieved, after which the resin was drained, and washed with DMF (4 × 5 mL) and dichloromethane (CH_2_Cl_2_) (5 × 5 mL).

After elongation, the fully-protected resin-bound peptide was subjected to acidolytic cleavage using TFA:*i*Pr_3_SiH:H_2_O (90:5:5 v/v/v) and the resulting suspension was gently agitated at room temperature for 2 hours. After filtration of resin, the cleavage cocktail was concentrated under vacuum and the residue was precipitated using diethyl ether. Upon centrifugation, the crude peptide obtained was subjected to reverse-phase HPLC purification (C18 column, gradient: 0 to 60% B over 50 min with 0.1% TFA, 15 mL/min) to afford neopentyl (nP) protected peptide. The nP protecting groups were removed by incubation of peptide at 50 °C in an aqueous buffer (6 M guanidine hydrochloride (Gdn•HCl), 0.1 M HEPES, pH 8) for 16 hours. After HPLC purification (C18 column, gradient: 0 to 75% B over 40 min with 0.1% ammonium hydroxide (NH_4_OH), 15 mL/min) the peptide was isolated in 3.2% overall yield (12 mg, 100 µmol scale).

### Purification of wild-type DARC

cDNA coding regions of ^WT^DARC^G42^ were cloned in pVL1393 vector with N-terminus HA-signal, FLAG-tag followed by M4 sequence and HRV 3C protease site. *Sf9* cells were infected with the baculovirus expressing ^WT^DARC^G42^ at a density of 1.8 million cells per ml and were cultured for 72 h in a shaking incubator at 135 rpm at 27 °C. Post 72 h, the cells were harvested by centrifugation at 5432×g in a Fiberlite F9-6 x 1000 LEX rotor (Thermo Scientific) for 10 minutes at 4 °C. The media was decanted, and the pellets were flash frozen in liquid nitrogen and stored at −80 °C till further use.

The receptor was purified using a similar protocol as discussed previously^9^. Briefly, cells expressing ^WT^DARC^G42^ were thawed and sequentially dounce-homogenized in hypotonic buffer (20 mM HEPES, pH 7.4, 10 mM MgCl_2_, 20 mM KCl, 1 mM PMSF and 2 mM benzamidine), and hypertonic buffer (20 mM HEPES, pH 7.4, 1 M NaCl, 10 mM MgCl_2_, 20 mM KCl, 1 mM PMSF and 2 mM benzamidine), followed by centrifugation at 48,000×g in a A27 - 8 x 50 Rotor (Thermo Scientific), for 20 min at 4 °C. Subsequently, the receptor was solubilised into detergent micelles using solubilisation buffer comprising 20 mM HEPES, pH 7.4, 450 mM NaCl, 1 mM PMSF and 2 mM benzamidine, in the presence of 0.1% (w/v) CHS, 0.5% (w/v) L-MNG, and 2 mM iodoacetamide for 2 h at 4 °C with constant tumbling. Following solubilisation, the lysate was diluted 3-fold in a dilution buffer comprising 20 mM HEPES, pH 7.4, 1 mM PMSF, 2 mM benzamidine and 2.5 mM CaCl_2_, followed by centrifugation at 30,000×g in a Fiberlite F14-6×250y rotor (Thermo Scientific) for 25 min at 4 °C. The supernatant was further passed through a 0.45 µm bottle-top filter and loaded onto M1 anti-FLAG resin in gravity-flow columns equilibrated with low salt buffer (LSB) comprising 20 mM HEPES, pH 7.4, 2 mM CaCl_2_, 0.1% (w/v) CHS and, 0.01% (w/v) L-MNG. The columns were then washed with 3 washes of LSB alternated with 2 washes of high salt buffer (20 mM HEPES, pH 7.4, 350 mM NaCl, 2 mM CaCl_2_ and 0.01% L-MNG). The bound receptor was eluted in a buffer comprising 20 mM HEPES, pH 7.4, 150 mM NaCl, 2 mM EDTA, 0.01% L-MNG, 250 μg/ml FLAG-peptide. Post-elution, free cysteines were blocked with 2 mM iodoacetamide, added twice in 10 min intervals followed by addition of 2 mM cysteine to quench excess iodoacetamide.

### Preparation of ^Sulfo^DARC

To purify N-terminally sulfated-DARC (^Sulfo^DARC), an *in-vitro* peptide ligation strategy mediated by bacterial sortase A enzyme was employed. ^WT^DARC^G42^ sequence was modified by insertion of a TEV cleavage site (ENLYFQ) after L^45^ in the N-terminus and the sortase ligation site (GGG), followed by the DARC sequence (^46^E-S^336^).The modified construct with TEV-site and sortase ligation site was named as DARC-sortase. To express the construct, *Sf9* cells were infected with the baculovirus expressing DARC-sortase at a density of 1.8 × 10^6^ cells/mL and grown in a suspension culture with a constant shaking at 135 rpm for 72 h at 27°C. The cells were harvested by centrifugation at 5432×g in a Fiberlite F9-6 x 1000 LEX rotor (Thermo Scientific) and the pellets were flash frozen in liquid nitrogen and stored at −80 °C till further use. The purification of DARC-sortase was performed similar to ^WT^DARC^G42^, however cysteine blocking was not performed. Post-elution, the native N-terminus of DARC-sortase was removed by TEV-protease. Specifically, the purified DARC-sortase receptor was incubated with TEV-protease (prepared in-house) at a ratio of 1:20 (TEV:receptor) at 25 °C for overnight. To remove the cleaved peptide, the reaction mix was then concentrated using a 100 kDa cutoff Vivaspin concentrator (Cytiva), and subjected to size exclusion chromatography (SEC) in a HiLoad^TM^ Superdex 200 column (Cytiva), pre-equilibrated with SEC buffer (20 mM HEPES, pH 7.4, 100 mM NaCl, 0.0001% (w/v) CHS, 0.00025% (w/v) GDN, 0.00075% (v/v) of L-MNG). The elution fractions corresponding to the receptor were pooled together.

To obtain ^Sulfo^DARC, sortaseA-mediated peptide ligation strategy was employed. The SEC purified N-terminal cleaved DARC-sortase was concentrated to 20-25 mg/mL using a 100 kDa cutoff Vivaspin concentrator (Cytiva) and incubated with 2.5 µM purified sortaseA (produced as outlined below) and 3-fold molar excess (of receptor) ^Sulfo^DARC peptide (harboring a sortaseA recognition sequence at its C-terminus, LPETG), in the presence of 5 mM CaCl_2_. The reaction mix was incubated at 25 °C for 3 h followed by quenching with 10 mM EDTA. The unligated peptide was separated from the ligated receptor (^Sulfo^DARC) by size exclusion chromatography (SEC) in a HiLoad Superdex 200 column (Cytiva), pre-equilibrated with the SEC buffer. The elution fractions corresponding to the ^Sulfo^DARC were pooled and flash frozen into liquid nitrogen and stored at −80 °C till further use.

### Purification of CCL7 and CXCL8

CCL7 was purified using a previously discussed protocol with slight modifications^9^. CCL7 and CXCL8 expression and purification were performed similarly unless stated otherwise. Briefly, *E.coli* BL21(DE3) cells were transformed with cDNA coding CCL7 and CXCL8 cloned in pGEMEX vector harboring 6X-His-tag followed by enterokinase cleavage site at its N-terminus. The cells were grown in terrific broth at 27 °C till O.D._600 nm_ 1.2 - 1.8, following which the culture was induced with freshly prepared 1 mM IPTG and cultured at 20 °C for 48 hours in a shaking incubator at 165 rpm. Post 48 h, the cell pellets were harvested by centrifugation at 5000 rpm for 20 minutes and flash-frozen in liquid nitrogen and stored at −80 °C till further use.

To purify the chemokines, the bacterial pellets were thawed and resuspended in the lysis buffer (30 mM MOPS, pH 7.2, 1 M NaCl, 10 mM imidazole, 0.5% (v/v) glycerol, 0.3% (v/v) triton X 100, 1 mM PMSF) and homogenised by constant stirring for 30 minutes at 4 °C, following which the cells were mechanically disrupted by sonication for 20 minutes with 15 s on and 30 s off pulse cycle. Subsequently, the lysed cells were centrifuged at 20,000 rpm for 20 minutes at 4 °C and the supernatant was filtered using 0.45 µm bottle-top filters. The lysate was then loaded onto Ni-NTA gravity flow columns pre-equilibrated with the equilibration buffer composed of 30 mM MOPS, pH 7.2, 1 M NaCl, 40 mM imidazole, 0.5% (v/v) glycerol at a flow-rate of 1 mL/min. The non-specific proteins were washed with the equilibration buffer and bound CCL7 and CXCL8 was eluted in 30 mM MOPS, pH 7.2, 1 M NaCl, 500 mM imidazole, 5% (v/v) glycerol. The eluted protein was dialysed against enterokinase cleavage buffer comprising 20 mM Tris-HCl, pH 8.0, 50 mM NaCl for 16 h at 4 °C. The N-terminus His-tag was removed by incubating dialysed CCL7 with enterokinase in the ratio of 1:200 (enterokinase: CCL7) in the presence of 10 mM CaCl_2_, for 24 h at 4 °C. The cleavage was confirmed by SDS-PAGE and the cleaved CCL7 and CXCL8 were separated from uncleaved by cation-exchange chromatography using ResourceS column and eluted with a linear gradient of NaCl. The elution fractions corresponding to cleaved CCL7 and CXCL8 were pooled together and dialysed against 20 mM HEPES, pH 7.4 and 150 mM NaCl Finally, the dialysed proteins were flash frozen in liquid nitrogen after adding 10% glycerol and stored at −80 °C till further use.

### Purification of Sortase

A modified penta-mutant sortase A (eSrtA) in pET29 was procured from Addgene (Addgene plasmid #<u>75144</u>) and purified using a previously described protocol^45^, with modifications. A single colony of BL21(DE3) cells transformed with the plasmid were inoculated into a 50 mL of LB media supplemented with 50 μg/mL kanamycin and incubated at 37 °C with constant shaking at 160 rpm, till O.D._600nm_ ∼0.6. This was then used to inoculate 1 L of a secondary culture in LB, which was grown, again at 37 °C with shaking at 160 rpm, till O.D_600nm_ reached ∼0.6. To induce protein expression, 100 μM IPTG was added, and the culture was grown with constant shaking at 160 rpm at 37 °C for 18 h. Pellets were harvested by centrifugation at 5432×g in a Fiberlite F9-6 x 1000 LEX rotor (Thermo Scientific) and the cell pellets were flash frozen in liquid nitrogen and stored at −80 °C till further use. For purification, pellets were resuspended in 5× pellet-volume of a lysis buffer composed of 50 mM Tris, pH 8.5, 300 mM NaCl and 1 mM MgCl_2_. Cells were ruptured by sonication for 20 min with 15 s on and 30 s off cycle. The resulting lysate was centrifuged at 48,000×g in a A27 - 8 x 50 Rotor (Thermo Scientific), for 20 min at 4 °C, to settle the debris. The resulting supernatant was clarified by passing through a 0.45 µm bottle-top filter and loaded onto a gravity-flow column containing Ni-NTA resin (Takara, cat no. 635662), at a flow rate of 1 mL/min. The resin was then washed using a wash buffer containing 50 mM Tris, pH 8.5, 150 mM NaCl, 0.1 mM MgCl_2_ and 30 mM imidazole, followed by elution using a buffer containing 50 mM Tris, pH 8.5, 150 mM NaCl, 0.1 mM MgCl_2_ and 250 mM imidazole. A part of this elutate was diluted in 4 x of MES buffer (pH 5.5) and clarified using anion exchange chromatography by loading onto a HiTrap^TM^ Q FF column (Cytiva), equilibrated with a buffer containing (50mM Tris-Cl, pH 8.0, 50mM NaCl). The peaks corresponding to the protein were pooled, and to remove salts, an overnight dialysis was performed using a buffer containing (20 mM HEPES pH 7.4, 150 mM NaCl). Dialysed protein was flash frozen in liquid nitrogen and stored in −80 °C until further use.

### NanoBRET binding assay

Ligand binding to ^WT^DARC^G42^ and mutated DARC was monitored by a NanoBRET binding assay^46,47^. Briefly, 5 × 10^6^ HEK293T cells were plated in 10-cm culture dishes and 24 h later transfected with vectors encoding either ^WT^DARC^G42^ or mutated DARC receptor, fused to N-terminal Nanoluciferase. Post 24 h transfection, cells were harvested and distributed into white 96-well plates at a density of 1 × 10^5^ cells per well. Cells were then incubated with AZDye 488-labelled chemokines for 2 h on ice (Protein Foundry) at concentrations ranging from 0.1 nM to 1 µM. Coelenterazine H (diluted 1:500) was then added and donor emission (450-80 nm BP filter) and acceptor emission (570-100 nm LP filter) were immediately measured on a PHERAstar FSX plate reader (BMG LABTECH). Concentration–response curves were fitted to the three-parameter Hill equation using an iterative, least-squares method (GraphPad Prism version 10.0.0). All curves were fitted to data points generated from the mean of at least three independent experiments.

### Cryo-EM sample preparation and data collection

For grid freezing of CXCL8-^Sulfo^DARC^G42^, CXCL8-^Non-sulfo^DARC^D42^ and CXCL8-^Sulfo^DARC^D42^ complexes, Quantifoil R1.2/1.3 Au 200-mesh grids were glow discharged at 20mA current for 60s glow and 10s hold using a PELCO easiGlow glow discharge system (*Ted Pella*). 3µL of the samples were applied onto glow-discharged grids, blotted for 5-6 s with blot-force of 0 at 4°C and 100 % humidity and plunged into liquid ethane using a Mark IV Vitrobot (Thermo Fisher Scientific).

For the CCL7-^WT^DARC^G42^, CXCL8-^WT^DARC^G42^ and CCL7-^Sulfo^DARC^G42^ complexes, 3 μL of the samples were dispensed onto glow discharged Quantifoil holey carbon grids (Au R1.2/1.3) using a PIB-10 glow-discharge (Vacuum Device Co., Ltd., https://www.shinkuu.co.jp/) and blotted using Ø55/20 mm, Grade 595 (Ted Pella, Inc.) filter paper. The grids were subsequently plunge-frozen in liquid ethane (−181°C) using a Vitrobot Mark IV maintained at 100% humidity and 4°C.

Data collection of the CXCL8-^Sulfo^DARC, CXCL8-^Non-sulfo^DARC^D42^ and CXCL8-^Sulfo^DARC^D42^ samples were performed on a 300 kV Titan Krios microscope (G4, Thermo Fisher Scientific) equipped with a K3 direct electron detector (Gatan) and a BioQuantum K3 energy filter. For the CCL7-^WT^DARC^G42^, CXCL8-^WT^DARC^G42^ and CCL7-^Sulfo^DARC^G42^ complexes, data collection was performed on a 300 kV Titan Krios microscope (G3i, Thermo Fisher Scientific) equipped with a K3 direct electron detector (Gatan) and a BioQuantum K3 energy filter. Multiframe movies were recorded in counting mode across a defocus range of −0.8 to −1.8 μm, at a pixel size of 0.86 Å/px using EPU software (Thermo Fisher Scientific) for the CXCL8-^Sulfo^DARC, CXCL8-DARC^D42^ and CXCL8-^Sulfo^DARC^D42^ samples, while movies against CCL7-DARC, CXCL8-DARC and CCL7-^Sulfo^DARC samples were recorded in counting mode across a defocus range of −0.8 to −1.6 μm, at a pixel size of 0.83 Å/px. Movies were collected with a total electron exposure of ∼55 e-/Å² for CXCL8-^Sulfo^DARC, CXCL8-^Non-sulfo^DARC^D42^ and CXCL8-^Sulfo^DARC^D42^ samples and 63.4 e-/Å² for the CCL7-^WT^DARC^G42^, CXCL8-^WT^DARC^G42^ and CCL7-^Sulfo^DARC^G42^ samples.

### Data processing

All datasets were processed in cryoSPARC v4 unless otherwise stated. Multiframe movies were imported and aligned with patch-based motion correction (multi), followed by contrast transfer function (CTF) estimation using patch CTF (multi). Processing of the DARC datasets were performed following a similar strategy as outlined in the accompanying pipeline (**Figure S1-2**). In brief, particles were picked using the integrated blob picker and extracted with an appropriate box size. Extracted particle stacks were subjected to multiple rounds of 2D classification to remove junk particles and contaminating ice. A clean particle stack from the 2D classes showing clear secondary structure features were subjected to ab-initio reconstruction. This step was followed by heterogeneous refinement, referenced to these ab initio models, further cleaning the particle stack. Particles corresponding to 3D classes exhibiting features of DARC-ligand complexes were subjected to non-uniform refinement, followed by local refinement of the ligand and receptor portions as needed. Final resolutions for all reconstructions were determined in cryoSPARC using gold-standard Fourier shell correlation (FSC) at 0.143 with half-maps.

Data collection, processing and model refinement statistics are provided in **Supplementary Table S1**. Local resolutions of all reconstructions were estimated using the Blocres algorithm, and maps were sharpened based on B-factor using the built-in sharpening tool in cryoSPARC v4.

### Model building and refinement

Initial models of DARC were derived from the coordinates of 8JPS (CCL7-DARC complex), whereas CXCL8 models were generated using SWISS-MODEL. These structures were docked into the B-factor sharpened map in UCSF Chimera^48^, followed by flexible fitting with the *all_atom_refine* module in COOT^49^. The fitted coordinates were subjected to iterative manual adjustments in COOT and refinement against the map using *phenix.real_space_refine* from the Phenix suite^50,51^. Final models showed excellent geometry, with the majority of residues in the most favoured regions of Ramachandran plots (**Supplementary Table S1**). All structural figures were prepared using Chimera and ChimeraX^52^.

