## Supplementary_Figures for "Molecular basis of promiscuous chemokine-engagement by the Duffy antigen receptor"

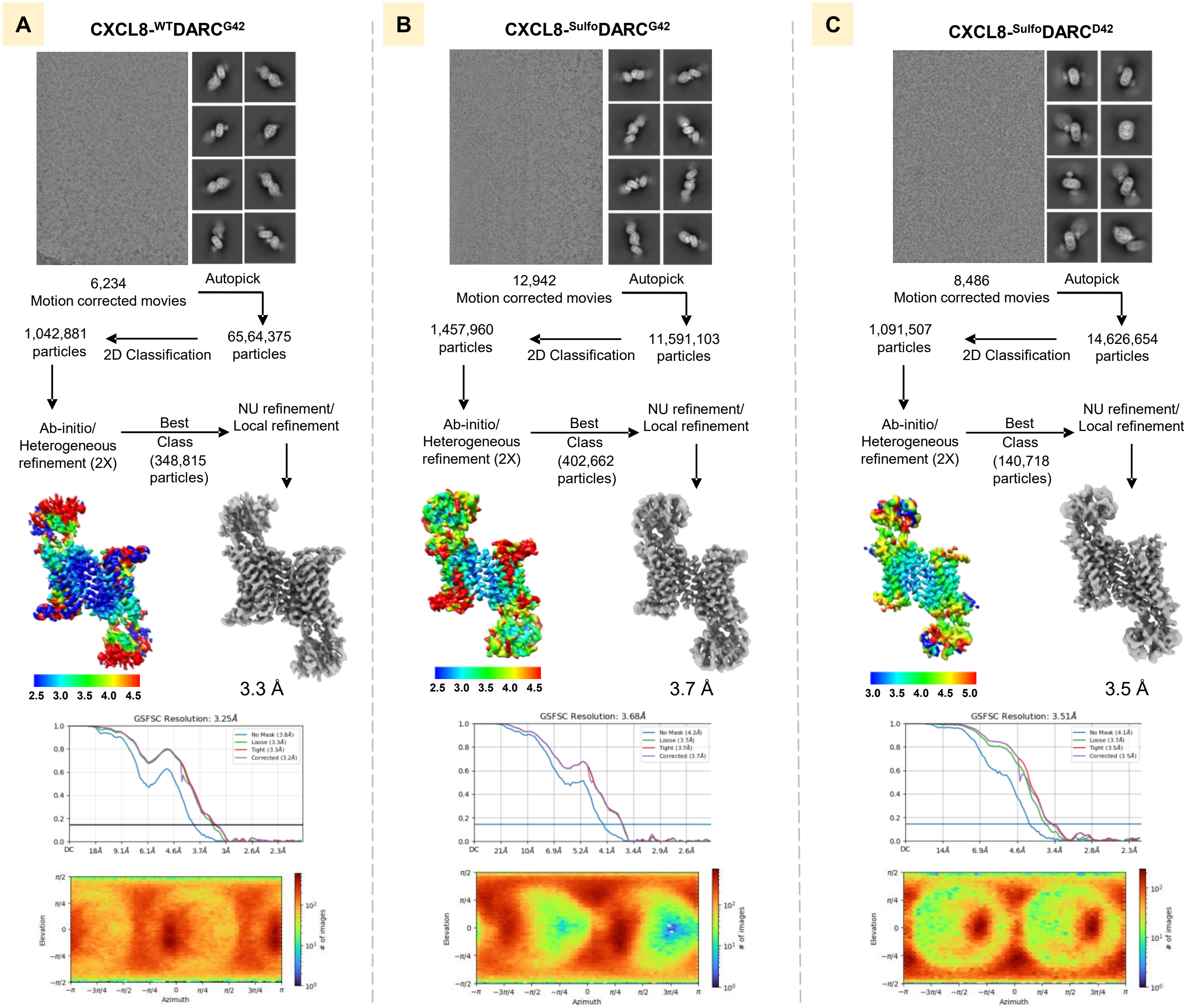


**Supplementary Figure S1: Workflow for cryo-EM data processing of DARC complexes.**
**(A-C)**, Representative cryo-EM micrograph, selected 2D class averages representing different orientations, schematic representation of cryo-EM data processing workflow, local resolution map of the 3D reconstruction, gold standard fourier shell correlation curve (GSFSC) at 0.143 threshold, and angular distribution of the particles against the final reconstruction of CXCL8-$^{WT}$DARC$^{G42}$, CXCL8-$^{Sulfo}$DARC$^{G42}$ and CXCL8-$^{Sulfo}$DARC$^{D42}$ complexes, respectively.

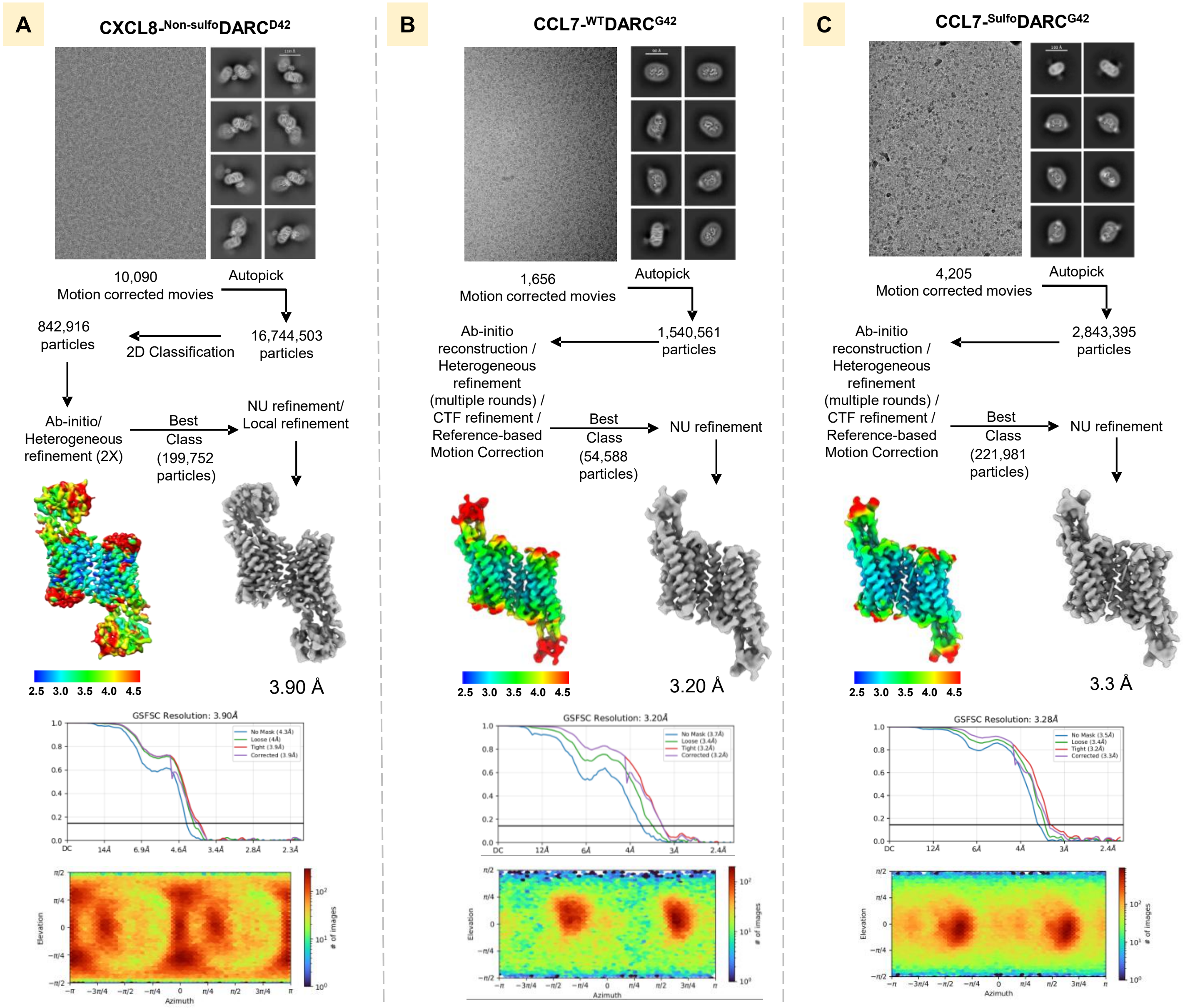


**Supplementary Figure S2: Workflow for cryo-EM data processing of DARC complexes.**
**(A-C)**, Representative cryo-EM micrograph, selected 2D class averages representing different orientations, schematic representation of cryo-EM data processing workflow, local resolution map of the 3D reconstruction, gold standard fourier shell correlation curve (GSFSC) at 0.143 threshold, and angular distribution of the particles against the final reconstruction of CXCL8-$^{Non\text{-}sulfo}$DARC$^{D42}$, CCL7-$^{WT}$DARC$^{G42}$ and CCL7-$^{Sulfo}$DARC$^{G42}$ complexes, respectively.

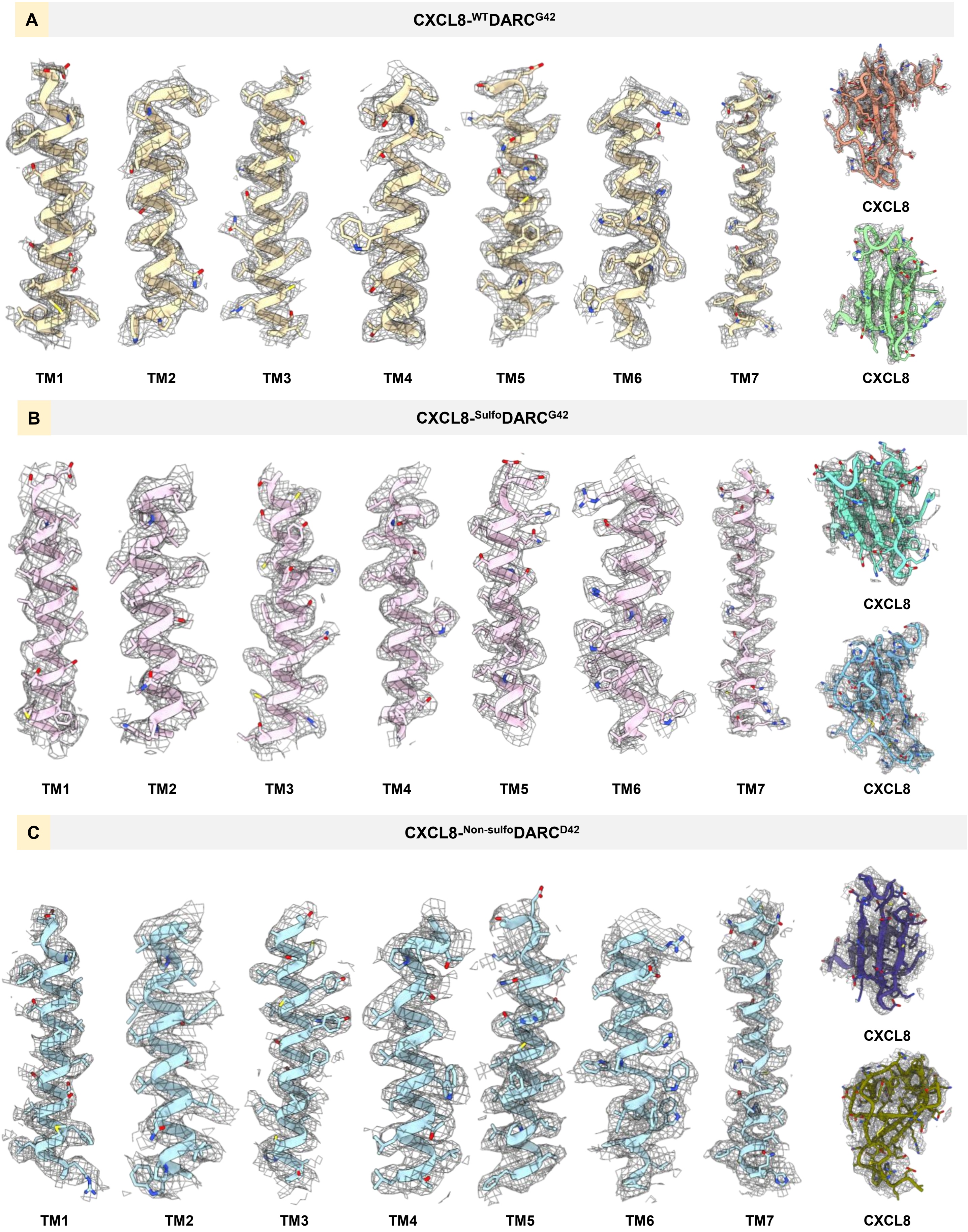


**Supplementary Figure S3. cryo-EM densities of the TM segments and ligands in DARC structures. (A-C)**, EM density maps of TM1-7, helix8 of CXCL8-$^{WT}$DARC$^{G42}$, CXCL8-$^{Sulfo}$DARC$^{G42}$, CXCL8-$^{Non\text{-}sulfo}$DARC$^{D42}$.

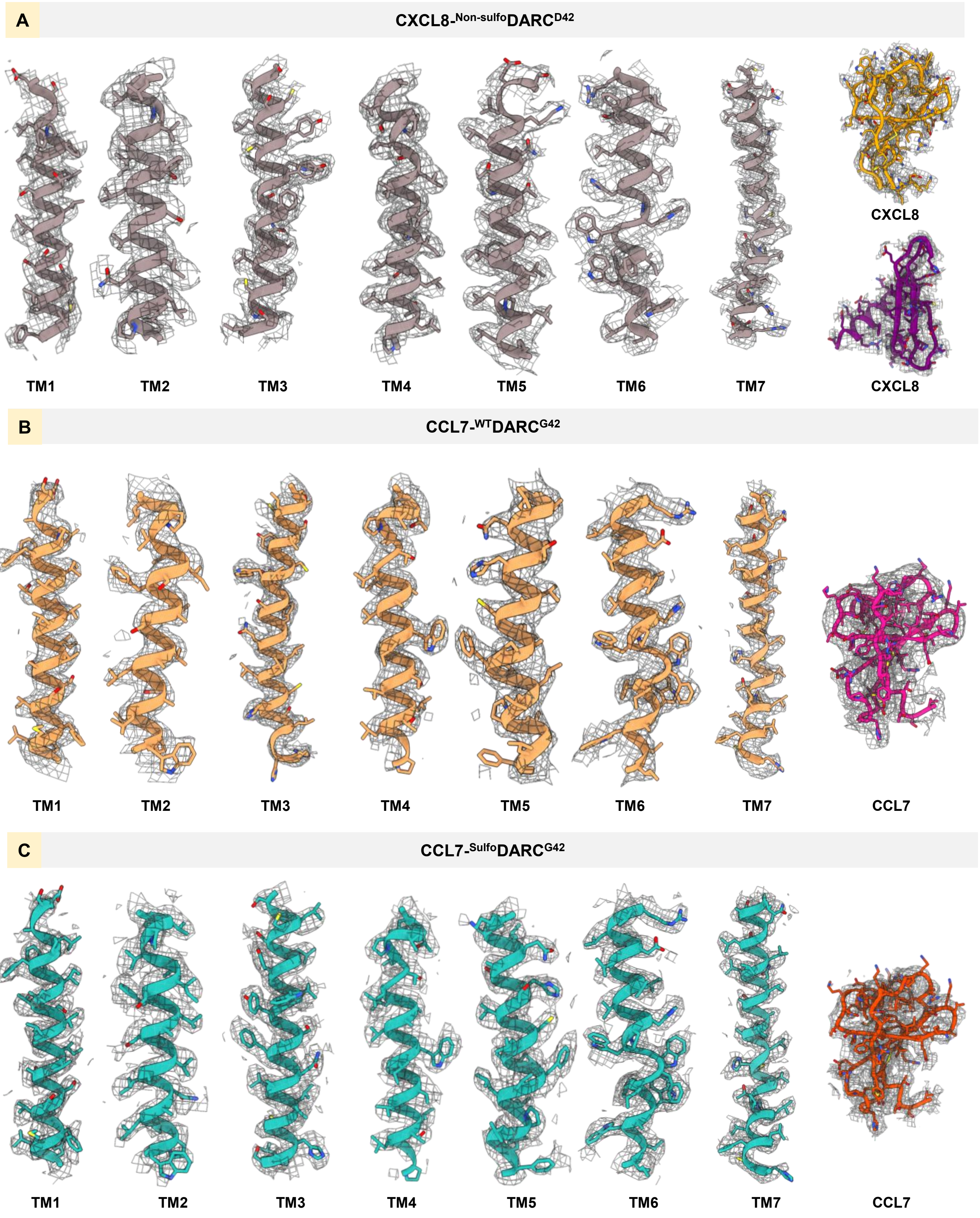


**Supplementary Figure S4. cryo-EM densities of the TM segments and ligands in DARC structures.** **(A-C)**, EM density maps of TM1-7, helix8 of CXCL8-$^{Sulfo}$DARC$^{D42}$, CCL7-$^{WT}$DARC$^{G42}$, CCL7-$^{Sulfo}$DARC$^{G42}$.

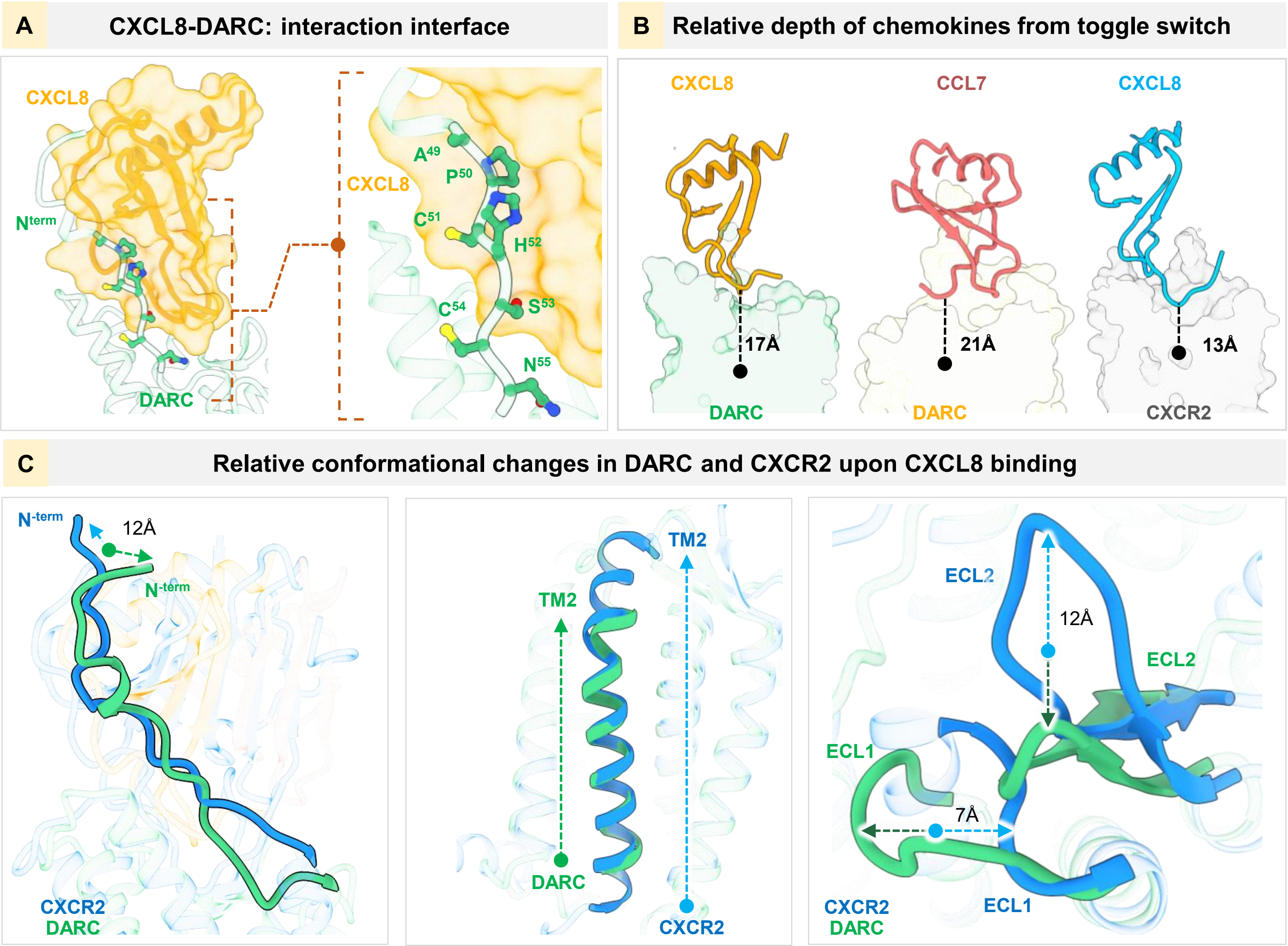


**Supplementary Figure S5. Structural insights into CXCL8-DARC interaction. (A)** Cartoon representation depicting the extensive interaction between the N-terminus of DARC with the CXCL8. **(B)** Relative depth of ligand penetration in DARC and CXCR2 structures, measured from conserved Trp$^{6.48}$ (PDB IDs: 21OZ for CXCL8-DARC, 21OX for CCL7-DARC, and 8XWN for CXCL8-CXCR2). **(C)** Conformational differences in DARC upon binding to CXCL8, compared to the CXCL8-CXCR2 structure (PDB ID: 8XWN).

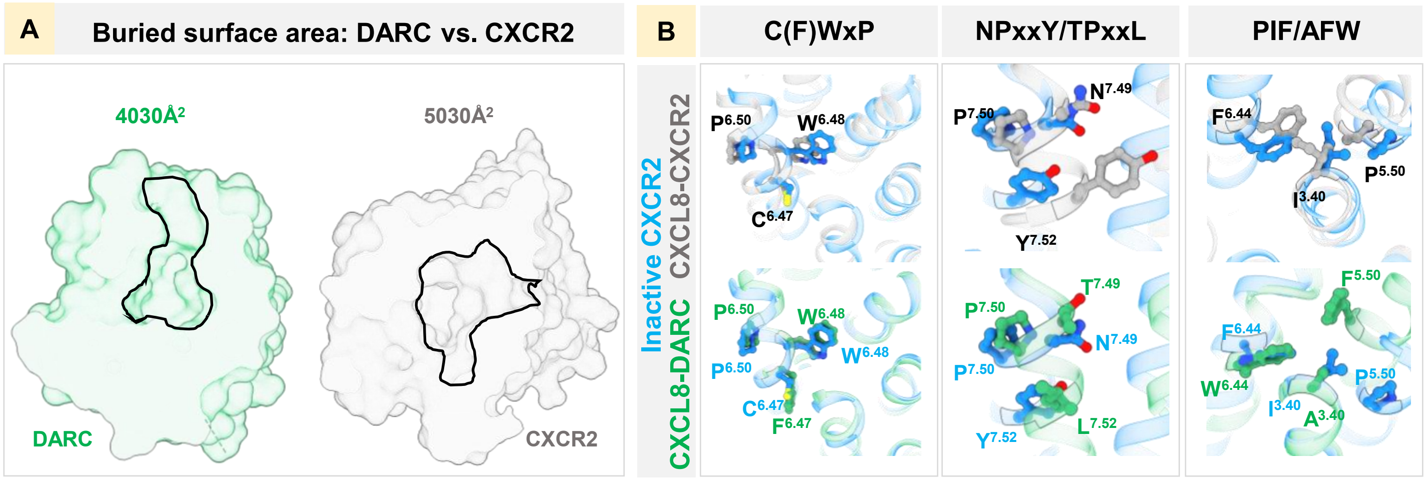


**Supplementary Figure S6. Structural comparison of CXCL8-bound DARC and CXCR2. (A)** A schematic of the buried surface area occupied by CXCL8 in DARC (PDB ID: 21OZ) vs. CXCR2 (PDB ID: 8XWN). **(B)** Structural changes in the microswitches between the inactive structure of CXCR2 (PDB ID: 6LFL) and CXCL8-bound CXCR2/DARC (PDB ID: 8XWN and 21OZ, respectively).

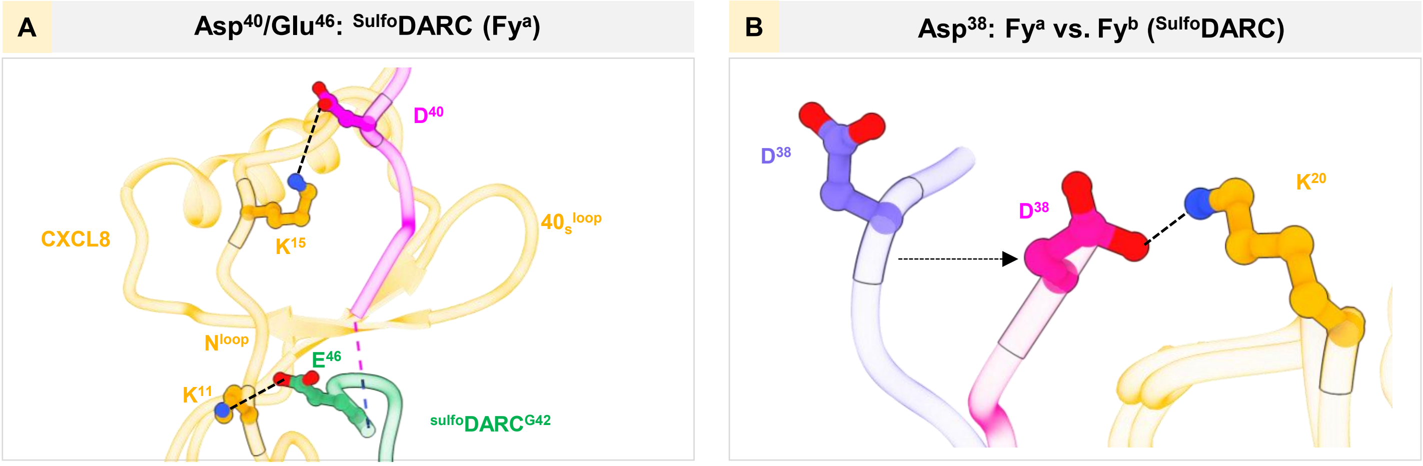


**Supplementary Figure S7. Structural comparison of the Fy$^{a}$ vs. Fy$^{b}$ variants of $^{Sulfo}$DARC. (A)** Asp$^{40}$ and Glu$^{46}$ interaction with N$^{loop}$ of CXCL8 residues are shown. **(B)** Shift of the Fy$^{b}$ N-terminus toward the core domain of CXCL8, highlighting the interaction between Asp$^{38}$ and Lys$^{20}$.
